# Multiomic Approach Characterises the Neuroprotective Role of Retromer in Regulating Lysosomal Health

**DOI:** 10.1101/2022.09.13.507260

**Authors:** James L. Daly, Chris M. Danson, Philip A. Lewis, Sara Riccardo, Lucio Di Filippo, Davide Cacchiarelli, Stephen J. Cross, Kate J. Heesom, Andrea Ballabio, James R. Edgar, Peter J. Cullen

## Abstract

Retromer controls cellular homeostasis through regulating integral membrane protein sorting and transport and by controlling late-stage maturation of the endo-lysosomal network. Retromer dysfunction, which is linked to neurodegenerative disorders including Parkinson’s and Alzheimer’s diseases, manifests in complex cellular phenotypes, though the precise nature of this dysfunction, and its relation to neurodegeneration, remain unclear. Here, we perform the first integrated multiomics approach to provide precise insight into the impact of Retromer dysfunction on endo-lysosomal health and homeostasis within a human neuroglioma cell model. We quantify profound changes to the lysosomal proteome, indicative of broad lysosomal dysfunction and inefficient autophagic lysosome reformation, coupled with a reconfigured cell surface proteome and secretome reflective of increased lysosomal exocytosis. Through this global proteomic approach and parallel transcriptomic analysis, we provide an unprecedented integrated view of Retromer function in regulating lysosomal homeostasis and emphasise its role in neuroprotection.

## INTRODUCTION

Due to the vast numbers of integral membrane proteins (and their associated proteins and lipids) that require efficient and timely transport through the endo-lysosomal network, the cellular consequences of network dysfunction are widespread. Commonly linked to neurodegenerative diseases through highly complex phenotypes, network dysfunction includes impaired synaptic transmission, accelerated secretion, and reduced lysosomal catabolism associated with the accumulation of damaged organelles and abnormal intracellular protein aggregates.

Retromer is a multiprotein complex that couples with accessory proteins to regulate the sequence-dependent sorting of hundreds of integral membrane proteins through the endo-lysosomal network, protecting them from lysosomal degradation^1, 2^. Moreover, Retromer plays a key role in regulating Rab7 nucleotide cycling^3–5^. Retromer deficiency has been observed in Alzheimer’s disease patient brain samples, where its depletion or dysfunction can trigger or accelerate amyloid-β (Aβ) and Tau pathologies ^6–10^. Subtle Retromer dysfunction is also associated with disease-causing mutations in Parkinson’s disease^11–16^ and selective deletion of a key Retromer gene in neurons causes an amyotrophic lateral sclerosis-like phenotype in mice^17^.

It is imperative to understand the global cellular consequences of Retromer depletion and/or dysfunction to fully appreciate and contextualise the increasing interest in Retromer as a potential therapeutic target for these diseases. Here, by employing an integrated proteomic approach, we provide an unprecedented view of the impact of Retromer dysfunction on endolysosomal homeostasis and health.

## RESULTS

### Retromer Depletion Induces Severe Morphological Changes to the Endo-lysosomal Network

We generated clonal knockout (KO) H4 neuroglioma cell lines targeting the core *VPS35* component of Retromer, and rescued these lines through stable re-expression of functional VPS35-GFP (**Extended Data Fig. 1A**)^18^. VPS35 depletion in H4 cells induced profound morphological changes to the endo-lysosomal network, far greater than observed in corresponding *VPS35* KO HeLa cells (**Figs. 1A-B, Extended Data Fig. 1B**). In VPS35 KO H4 cells, transmission electron microscopy (TEM) revealed dramatically enlarged hybrid endo-lysosomal compartments, up to 10 µm in diameter, loaded with undigested membranous intraluminal material (**Fig. 1C**). These structures are reminiscent of those observed in Alzheimer’s, Parkinson’s and Lewy Body disease patients^19–21^. BSA-gold uptake assays unambiguously defined the enlarged hybrid compartments as endocytic in nature (**Extended Data Figs. 1C-D**). The pool of small, dense-core lysosomes observed in wild-type cells was depleted in VPS35 KO cells, suggesting that the swollen VPS35 KO compartments arrive from endosome-lysosome fusion events, followed by subsequent failure to clear the luminal content and reform lysosomes from the hybrid compartment. Importantly, VPS35-GFP re-expression facilitated the clearance of this accumulated material to rescue endo-lysosomal morphology and the population of dense-core lysosomes (**Fig. 1C**). Epidermal growth factor (EGF) uptake assays confirmed that aberrant VPS35 KO lysosomes exhibited a limited degradative capacity (**Extended Data Figs. 1E-G**).

**Figure 1.**
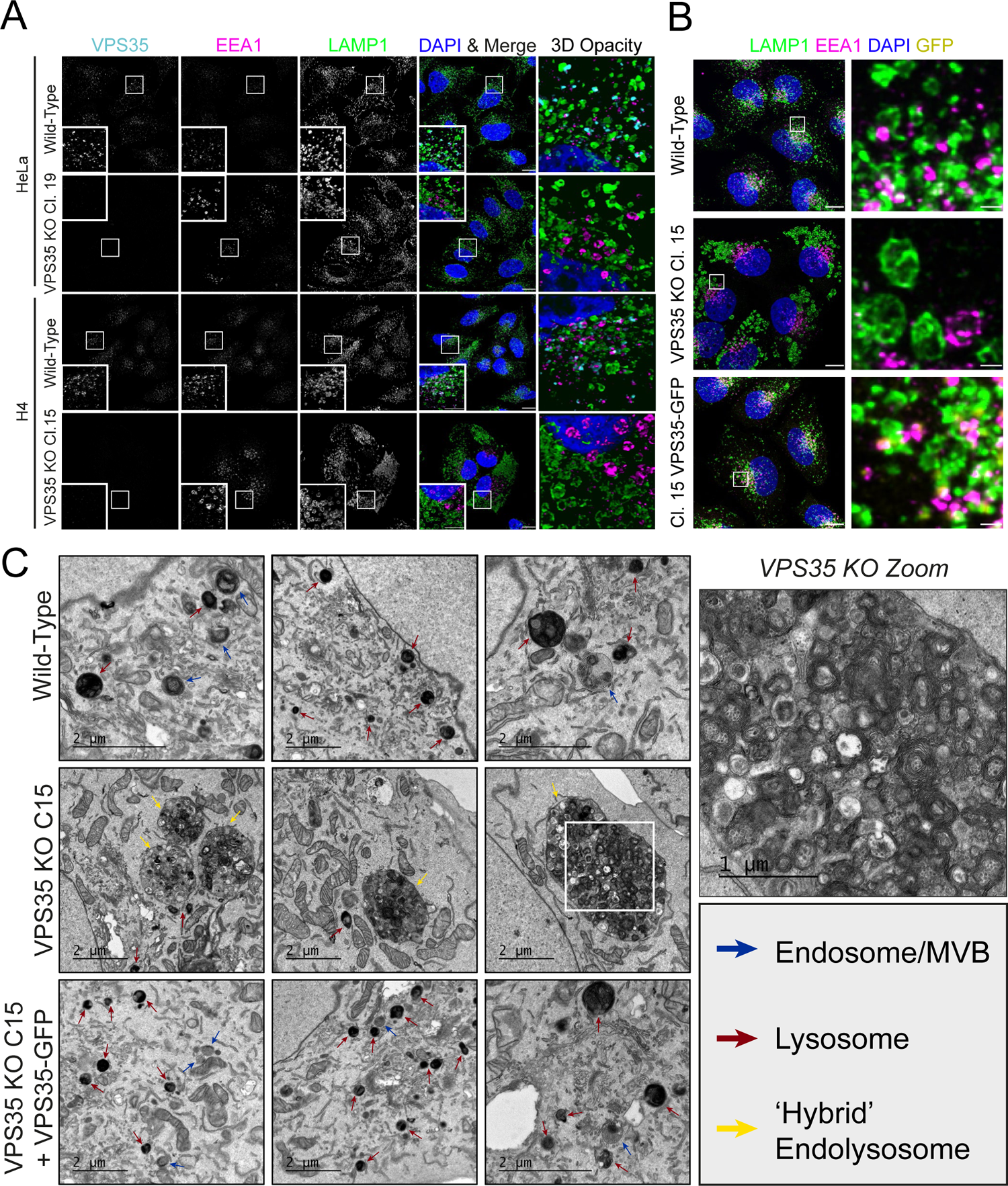
Retromer Depletion Imposes Severe Morphological Changes on the Endolysosomal Network in H4 Neuroglioma Cells. **(A)** The severity of endolysosomal swelling is enhanced in H4 VPS35 KO relative to HeLa cells. H4 and HeLa were transfected with CRISPR-Cas9 plasmids targeting VPS35, prior to clonal KO lines being fixed and immuno-stained for VPS35, EEA1, LAMP1 and DAPI. Scale bars: 20 µm and 5 µm in zoomed panels. **(B)** Morphological changes to the endolysosomal network in VPS35 KO are rescued by re-expression of VPS35-GFP. Wild-type, VPS35 KO Cl.15 and Cl.15 + VPS35-GFP H4 cells were fixed and immuno-stained for LAMP1, EEA1 and DAPI. Scale bars: 20 µm and 2 µm in zoomed panels. **(C)** VPS35 KO H4 exhibit enlarged hybrid endolysosomal compartments in which undigested materials accumulate. Transmission electron micrographs of endolysosomal compartments in cell lines, blue arrows denote endosomes/multivesicular bodies (MVBs), red = lysosomes and yellow = hybrid endo-lysosomes. Magnified panel depicts enlarged hybrid compartment in VPS35 KO. Scale bars: 2 µm and 1 µm in zoomed panel.

### LysoIP Proteomics Reveals a Fingerprint of Lysosomal Dysfunction in VPS35 KO Cells

To unbiasedly define the altered lysosome morphology in VPS35 KO H4 cells we coupled a lysosome immunoprecipitation (LysoIP) methodology with quantitative proteomics (**Fig 2A**)^22^. We transduced wild-type, VPS35 KO and VPS35-GFP-expressing H4 with transmembrane protein 192 (TMEM192), C-terminally flanked by three tandem HA epitopes (**Extended Data Fig. 2A**). Immunostaining of transduced cells with anti-HA and LAMP1 antibodies confirmed the specificity of TMEM192-x3-HA to label lysosomes (**Fig. 2B**). Immunoblotting of anti-HA LysoIPs showed a strong enrichment for the lysosomal marker LAMP1 that appeared highly consistent between wild-type, VPS35 KO and VPS35-GFP rescue-derived lysosomes (**Fig 2C**). From isobaric tandem mass tagging (TMT) and LC-MS/MS quantification we obtained a data set from wild type cells highly enriched for lysosomal proteins (**Extended Data Figs. 2B-C**), which contained 709 of the 828 proteins previously described as associated with lysosomes in HEK293T cells (**Fig. 2D**)^22^.

**Figure 2.**
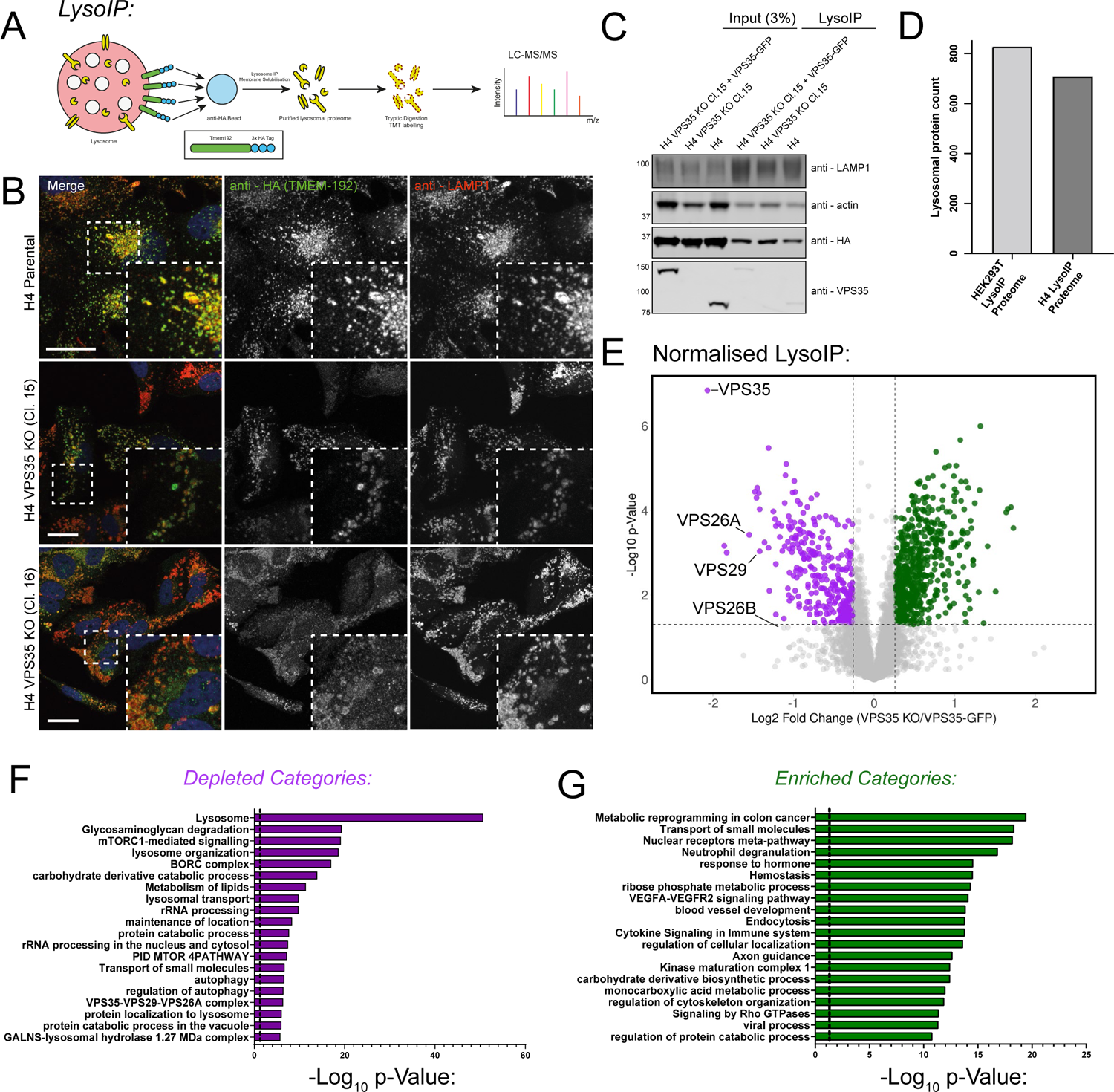
Development of organelle restricted proteomics to characterise VPS35 KO lysosomes. **(A)** Schematic depicting LysoIP methodology coupled to TMT-based quantitative proteomics. **(B)** TMEM192–x3HA labels lysosomes with a high degree of specificity. Representative confocal images of H4 cell lines transduced to express TMEM192-x3 HA prior to fixation and immuno-staining for HA, LAMP1 and DAPI. Scale bars: 20 µm. **(B)** LysoIP efficiently precipitates lysosomes from wild-type, VPS35 KO and VPS35-GFP rescue cell lines. LysoIP was performed on indicated cell lines prior to immuno-blotting with anti-LAMP1 (to assess lysosome enrichment), β-actin, HA and VPS35. **(C)** The lysosomal proteome of H4 exhibits a high degree of overlap with the published proteome of HEK293T-derived lysosomes. **(E)** A cohort of proteins are significantly relatively enriched (477 proteins) or depleted (246 proteins) from VPS35 KO lysosomes compared to both wild-type and rescue cells. The VPS35 KO/VPS35-GFP abundance ratio is displayed as a volcano plot. Data were normalised relative to total protein count and used to generate a volcano plot from n=7 independent experiments (clone 15 (n=3) and clone16 (n=4)), presented as ratio of Log_2_ VPS35 KO / KO + VPS35 GFP vs -Log10 p-value (thresholds set at p=0.05 and fold Log_2_ change +/- 0.26). **(F, G)** Cohorts of depleted (F) or enriched (G) proteins in VPS35 KO lysosomes converge into functional groupings. Gene ontology analyses of significantly enriched or depleted proteins (log_2_ fold change ± 0.26, p <0.05), describing depleted and enriched functional categories plotted against significance.

We generated proteomic LysoIP datasets from wild-type, VPS35 KO and rescue cell lines. VPS35 KO H4 cells exhibited a dramatic increase in protein abundances following LysoIP, reflective of the increased lysosomal number and size observed by microscopy, and consistent with the known activation of the master lysosomal biogenesis regulator transcription factor EB (TFEB) upon Retromer dysfunction in various cell types^3, 23^ (**Extended Data Fig. 3A, Table 1**).Data were therefore normalised based on total protein count to provide relative changes to the lysosomal proteome (**Fig. 2E, Table 2**). Comparison of LysoIP proteomics from VPS35 KO H4 cells relative to wild-type and VPS35-GFP rescues, revealed significantly depleted (246 proteins) and enriched (477 proteins) proteins (log_2_ fold change ± 0.26, p <0.05). Gene ontology analysis revealed an overall loss in lysosomal identity and an increased abundance of various pathways including metabolic reprogramming and small molecule transport (**Figs. 2F-G, Table 3**).

**Figure 3.**
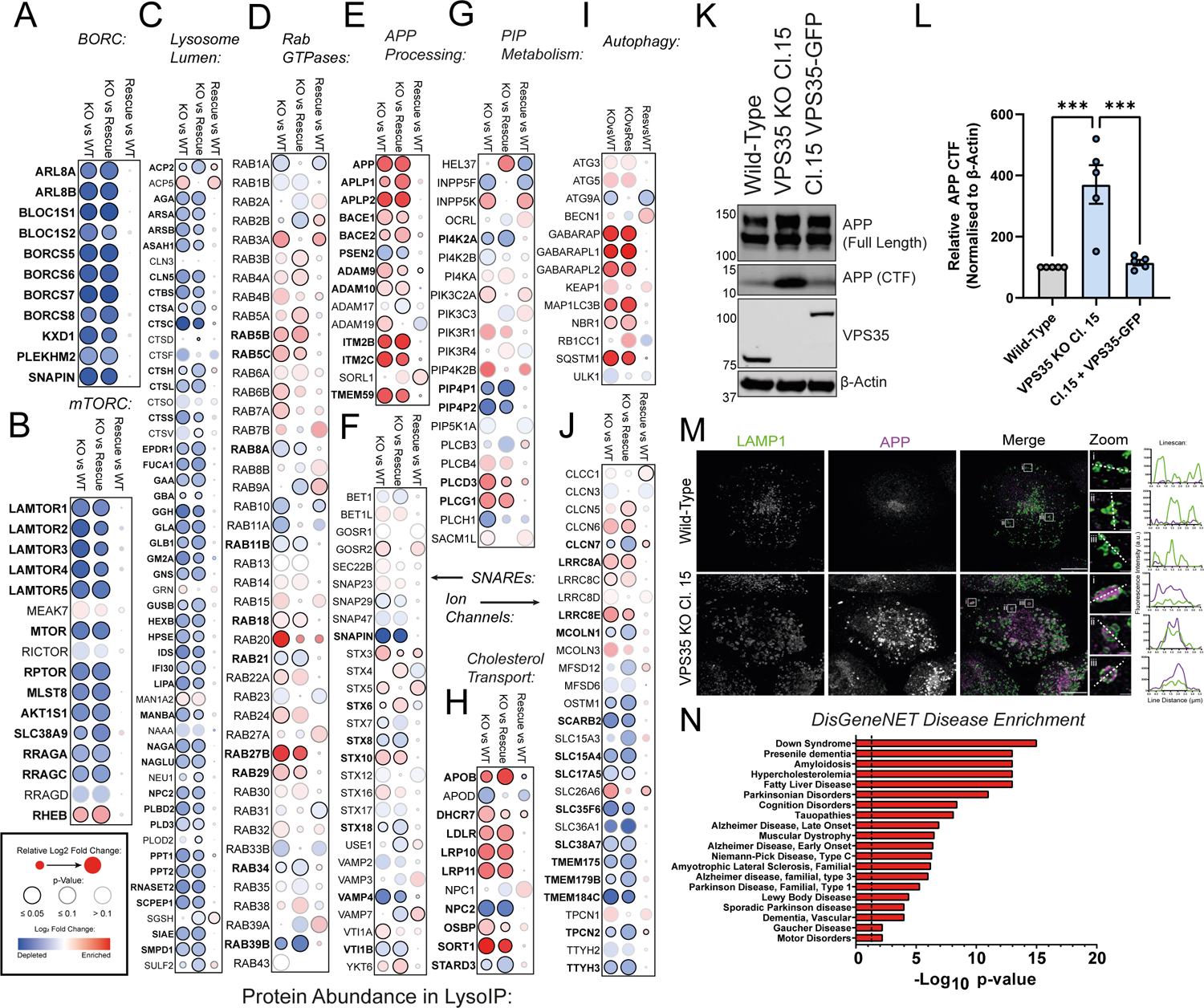
LysoIP Proteomics Reveals a Fingerprint of Lysosomal Dysfunction in VPS35 KO Cells. (**A-J**) A range of functional networks and protein families were relatively depleted or enriched in lysosomes derived from VPS35 KO H4 cells relative to wild-type and VPS35-GFP-expressing rescue cells. Dot-plots representing log_2_ fold change and p-value in quantified abundances of: (A) BORC; (B) mTORC1 complexes; (C) lysosomal lumen proteins; (D) Rab GTPases; (E) APP processing proteins; (F) vesicle SNAREs; (G) PIP metabolism; (H) cholesterol transport; (I) autophagy; and (J) lysosomal solute channels. (**K**) Proteolytic processing of APP is enhanced in VPS35 KO H4 cells. Cell lysates from the indicated cell lines were immuno-blotted using anti–APP (full length and CTF), VPS35 and β-actin antibodies. APP-CTF levels in VPS35 KO clone were rescued upon re-introduction of VPS35-GFP. (L) Quantification of APP-CTF signal intensity (n=5 independent experiments), means ± SEM, ordinary one-way ANOVA with Tukey’s multiple comparisons tests, wild-type vs VPS35 KO p = 0.0006, VPS35 KO vs VPS35-GFP p = 0.0009, wild-type vs VPS35-GFP p = 0.9540. **(M)** APP accumulates in LAMP1-positive compartments in VPS35 KO cells. Wild-type and VPS35 KO H4 were fixed and immuno-stained for LAMP1 and APP and linescan analysis was used to demonstrate colocalisation. Scale bar = 20 µm and 1 µm in zoomed panels. **(N)** Proteins enriched in the VPS35 KO LysoIP dataset are associated with neurodegenerative disease. Enrichment of selected DisGeneNET disease categories represented by significantly enriched proteins in the VPS35 KO LysoIP dataset (log_2_ fold change ± 0.26, p <0.05) relative to both wild-type and VPS35-GFP-expressing control conditions.

A range of protein-protein interaction networks were significantly depleted and enriched in the VPS35 KO LysoIP dataset (**Extended Data Figs. 3B-C**). All components of the BORC complex were depleted from VPS35 KO lysosomes and rescued by VPS35-GFP re-expression (**Fig. 3A**). The BORC complex positions lysosomes by coupling to kinesin-mediated microtubule transport via the adaptor protein Arl8, which was also depleted from VPS35 KO lysosomes^24^. Lysosomal recruitment of BORC is regulated by mTORC1/Ragulator. Indeed, mTOR and associated components including all Ragulator subunits LAMTOR1-5 and RagA/C GTPases were depleted in VPS35 KO lysosomes (**Fig. 3B, Extended Data Fig. 3D**). Rheb, an amino acid-responsive activator of mTORC1^25^, was the only component of the mTORC1 machinery to be significantly enriched in VPS35 KO lysosomes (**Fig. 3B**).

A wide cohort of luminal proteins, a large proportion of which are hydrolytic enzymes, were significantly depleted in VPS35 KO cells, including proteases, lipases, nucleases, and glycosidases including β-galactosidase (GLB1), which has previously shown to exhibit reduced activity in Retromer-depleted HeLa cells^26^ (**Fig. 3C**). Notably, the lysosomal acid glucosylceramidase GBA, which is genetically linked to Gaucher’s disease and Parkinson’s disease, was significantly depleted in VPS35 KO lysosomes^27^, as were a wider cohort of enzymes associated with lysosomal storage diseases^28^.

Rab GTPases are crucial regulators of membrane identity and transport^29^. Rab5a and Rab5b were significantly enriched in lysosomes from VPS35 KO H4 cells, reflective of increased mixing between early and late endosomes and defective resolution of these compartments (**Fig. 3D**). Rab7a was also significantly enriched in VPS35 KO cells compared to wild-type controls, as was Rab29, with the strongest enrichment being Rab27b, a late endosomal GTPase that regulates lysosomal exocytosis. Rab GTPases involved in late endosomal fusion with phagosomes and autophagosomes were dysregulated, including Rab20, Rab21 and Rab34^30–32^. Rab39b, which has been reported to regulate alpha-synuclein accumulation and has loss-of-function mutations in X-linked Parkinson’s disease, was strongly depleted in VPS35 KO lysosomes^33, 34^ (**Fig. 3D**).

Within the data we also identified: an enrichment of proteins involved in APP processing and metabolism, including an accumulation of APP in VPS35 KO lysosomes (**Fig. 3E**); depletion of the late endosomal SNARE proteins STX8 and VTI1B (**Fig. 3F**); dysregulation of phosphatidylinositol-4-phosphate (PI(4)P) metabolism, including depletion of PIP4P1 (TMEM55B), a regulator of perinuclear lysosomal transport and v-ATPase assembly^35, 36^ (**Fig. 3G**); perturbation of cholesterol influx and egress from lysosomes, which has been linked strongly with late onset, atypical cognitive decline^37^ (**Fig. 3H**); enrichment of autophagy markers (**Fig. 3I**); and alterations to lysosomal solute channels that regulate transmembrane transport to maintain lysosomal homeostasis, including CLCN7, a key Cl^-^/H^+^ antiporter that balances charge within the lysosomal lumen^38^, and TMEM175, recently linked to Parkinson’s disease^39^ (**Fig. 3J**). Taken together, these data highlight the scale of lysosomal dysfunction induced by Retromer depletion, many of which are linked to neurodegenerative disease.

We focussed on the enrichment of APP, which shuttles between the *trans*-Golgi network (TGN), endosomal network and plasma membrane under steady-state conditions and can undergo multiple proteolytic cleavage steps performed by α-, β-, and γ-secretase enzymes^40^. APP mutations that alter its proteolytic processing are causally linked to Alzheimer’s disease through the generation of neurotoxic amyloid aggregates ^40^. We noticed that VPS35 KO H4 cells demonstrate an abnormal accumulation of cleaved APP in the whole cell lysate, measured as a low molecular weight C-terminal fragment resulting from α- or β-secretase cleavage events (**Figs. 3K-L**). A similar accumulation of APP CTFs was recently reported in Vps35-depleted mouse neurons^41^. Levels of full-length APP were unaffected **(Fig. S3E)**. In wild-type and VPS35-GFP-expressing cells, APP predominantly localises within the perinuclear region, mainly colocalising with TGN46 and displaying minimal overlap with LAMP1 (**Extended Data Fig. 3F**). In contrast, VPS35 KO cells demonstrated a striking accumulation of APP on the limiting membrane or within LAMP1-positive compartments (**Fig. 3M** and **Extended Data Fig. 3F**). APP misprocessing and accumulation within the endo-lysosomal network is a crucial hallmark of Alzheimer’s disease, and is sufficient to induce lysosomal dysfunction^42^. Interestingly, DisGeneNet categories associated with significantly enriched proteins on VPS35 KO lysosomes included ‘Down Syndrome’, ‘Presenile Dementia’, ‘Amyloidosis’, ‘Alzheimer Disease’, ‘Parkinson Disease’, amongst others, indicating that the proteomic changes observed in these cells correlate with established phenotypes of neurodegeneration (**Fig. 3N, Table 4**).

### Correlative Proteomic Analyses Reveal a Signature of Lysosomal Exocytosis in VPS35 KO Cells

Retromer dysfunction induces a widespread reduction in the cell surface expression of integral membrane proteins as they become sequestered within internal endolysosomal compartments^43^. By coupling surface restricted biotinylation and LysoIP methodologies with quantitative proteomics, we attained a global, unbiased overview of how Retromer depletion triggered shifts in integral membrane protein abundances at the cell surface and lysosome, respectively (**Figs. 4A, Extended Data Figs. 4A-B, Table 2**). We observed a cohort of integral membrane proteins that were depleted from the cell surface and became enriched in the lysosome in VPS35 KO cells, including known Retromer cargoes such as GLUT1 (SLC2A1), STEAP3, SEMA4B and SEMA4C, NOTCH1 and NOTCH2, NETO2, KIDINS220, the neutral amino acid transporters SLC1A4 and SLC1A5, and the copper transporters ATP7A and SLC31A1^43–46^ (**Fig. 4B**). We validated GLUT1 and ATP7A re-routing from the cell surface into TMEM192-3xHA-positive lysosomes by Western blotting and immunostaining (**Fig. 4C, Extended Data Fig. 4C**) A number of these cargoes, for example SEMA4B/C, NOTCH1/2, NETO2 and KIDINS220, regulate essential neuronal pathways such as axonal guidance^47, 48^, and synaptic transmission^49–51^, and are associated with neuronal disease and disorders^52, 53^. These data underscore the protective role of Retromer in regulating the integrity of the plasma membrane proteome alongside maintaining lysosomal function.

**Figure 4.**
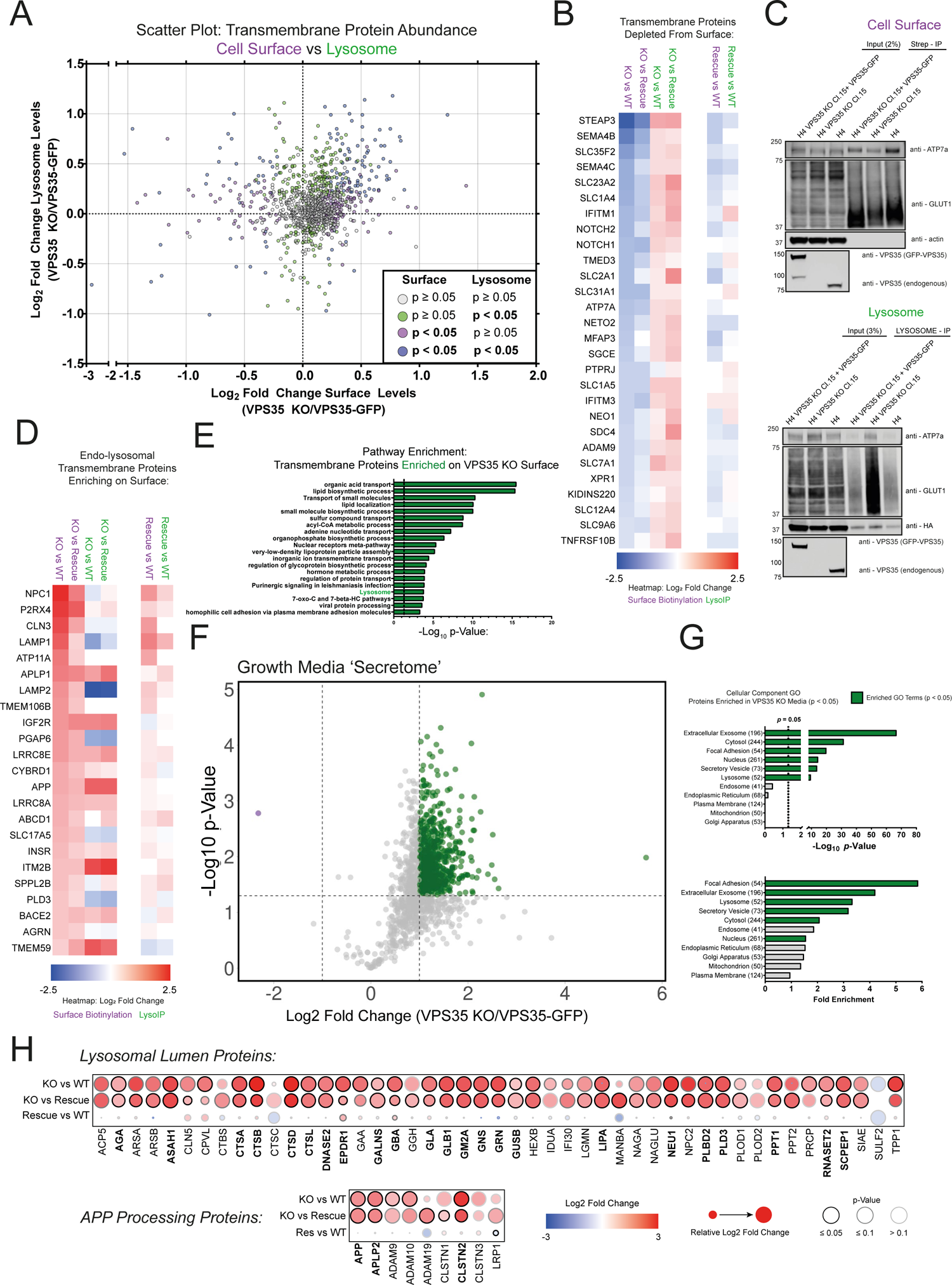
The cell surface proteome is re-modelled in VPS35 KO H4 cells. **(A)** Correlative analysis between cell surface and lysosomal proteome reveals bi-directional re-routing of transmembrane proteins. Scatter plot of VPS35 KO/VPS35-GFP transmembrane protein abundances in the cell surface proteome (x-axis) versus the LysoIP proteome (y-axis). Datapoints are coloured based on p-value scores in each experiment as denoted in the key. **(B)** Transmembrane proteins depleted from the cell surface are enriched in lysosomes of VPS35 KO cells. Heatmap depicting the loss of transmembrane proteins from the cell surface proteome (purple columns) with concomitant enrichment in the LysoIP proteome (green columns) in VPS35 KO cells relative to wild-type and VPS35-GFP rescue controls. **(C)** Perturbed endosome to plasma membrane recycling correlates with increased abundance of cell surface receptors in lysosomes. Cell surface and lysosomal proteomes were subjected to immuno-blotting with antibodies recognising defined Retromer cargoes (ATP7a and GLUT1). **(D)** Transmembrane lysosomal proteins enriched at cell surface VPS35 KO cells. Heatmap depicting the enrichment of transmembrane proteins in the cell surface proteome (purple columns) and corresponding abundance in the LysoIP proteome (green columns) in VPS35 KO cells relative to wild-type and VPS35-GFP rescue controls. **(E)** Analysis of significantly enriched pathways in the VPS35 KO transmembrane cell surface proteome relative to wild-type and VPS35-GFP rescue controls. **(F)** Volcano plot of VPS35 KO/VPS35-GFP protein abundances in the growth media ‘secretome’. 325 proteins were significantly enriched in the growth media of VPS35 KO cells compared to both wild-type and rescue sample (Log_2_ fold change > 1, p < 0.05). (**G**) Gene ontology analysis of cellular components enriched in the VPS35 KO ‘secretome’. Statistically significant categories (p < 0.05) are displayed in green. (**H**) Dot-plot predicting lysosomal luminal protein and APP processing protein abundances in the VPS35 KO ‘secretome’ relative to wild-type and VPS35-GFP rescue control samples. Log_2_ fold change, relative change and p-value score are depicted by dot colour, size and outline respectively, as depicted in the legend.

Interestingly, a cohort of lysosomal integral membrane proteins became enriched on the cell surface in VPS35 KO H4 cells, including the lysosomal glycocalyx LAMP1 and LAMP2, the cholesterol transporter NPC1, the Juvenile Neuronal Ceroid Lipofuscinosis associated CLN3, CI-MPR (IGF2R), and a cluster of APP processing-related proteins including APP, the β-secretase BACE2, the APP-like protein and β-secretase substrate APLP1^54^, and APP-binding protein ITM2B^55, 56^ (**Fig. 4D**). Specifically, the C-terminal fragment of APP was found to enrich at the cell surface in VPS35 KO cells (**Extended Data Figs. 4C-D**). Gene ontology analysis reflected these changes, including depletion of pathways related to cell morphogenesis, adhesion, synapse organisation and transmembrane transport (**Extended Data Fig.4E),** which correlate with enrichment in the lysosome (**Extended Data Fig.4F**), and a concomitant increase in metabolic pathways and lysosomal proteins at the cell surface, amongst others (**Fig. 4E, Table 3**).

Enrichment of lysosomal proteins at the cell surface could reflect increased lysosomal exocytosis, a process considered to be compensatory for lysosomal stress that may mediate cell-to-cell transfer of pathogenic aggregates such as α-synuclein and APP fragments^57–60^. A cell culture ‘secretome’ revealed a dramatic increase in proteins in the growth media of VPS35 KO H4 cells that was rescued by VPS35-GFP re-expression (**Fig. 4F, Table 1**). Gene ontology analysis revealed a significant enrichment of ‘focal adhesion’, ‘extracellular exosome’, ‘lysosome’ and ‘secretory vesicle’ cellular component categories **(Fig, 4G, Table 5**). Moreover, autophagic cargo receptor Sequestome-1 (SQSTM1) was prominently enriched in the secretome of VPS35 KO cells, suggestive of the release of autophagic material (**Table 1**). A wide cohort of luminal lysosomal proteins, which were relatively depleted in the VPS35 KO LysoIP dataset (**Fig. 3C**), were significantly enriched in the VPS35 KO secretome (**Fig. 4H**). Among these, some have been reported to undergo secretion from the biosynthetic pathway upon Retromer depletion^61–63^. However, we observed that CTSD colocalised completely with LAMP1 and displayed no evidence of accumulation within the biosynthetic pathway (**Extended Data Fig. 4G**). We therefore posit that in addition to biosynthetic pathway ‘leakage’, the delivery of lysosomal enzymes can be maintained in VPS35 KO cells, and their extracellular release may arise from lysosomal exocytosis, with CTSD being predominantly detected in a precursor state due to ineffective pH-dependent proteolytic activation in the endo-lysosomal network.

We also noticed an enrichment of APP in the secretome of VPS35 KO cells, alongside related substrates of the β- and γ-secretase enzymes APLP2 and CLSTN2, respectively^54, 64^ (**Fig. 4I**). Impaired APP processing and increased extracellular release of APP and Aβ fragments have been reported upon VPS35 depletion^9, 65^. Moreover, APLP2 was the most significantly enriched protein in the cerebrospinal fluid of Vps35 KO mice^10^. APP enrichment within the VPS35 KO secretome may therefore be indicative of altered amyloidogenic processing and release from the cell surface.

Taken together, these data indicate an increased release of lysosomal contents from the cell surface in response to the profound lysosomal stress observed in VPS35 KO H4 cells. This emphasises the neuroprotective role of Retromer in safeguarding against lysosomal dysfunction and extracellular release of cytotoxic contents such as APP cleavage products. These findings raise the possibility that the detection of APP, APLP2 or other components of the VPS35 KO secretome could be diagnostic tools for identifying Retromer dysfunction in cellular and model organism systems.

### RNA Sequencing of VPS35 KO H4 Cells Reveals Transcriptional Reconfigurations

Retromer dysfunction is known to induce TFEB activation^3, 23^. To directly investigate whether changes in the lysosomal proteome in VPS35 KO H4 cells were due to a coordinated transcriptional response through, for example, the co-ordinated lysosomal expression and regulation (CLEAR) network, we performed RNA sequencing (RNA-seq) of wild-type, VPS35 KO and VPS35-GFP H4 cells (**Extended Data Figs. 5A-B, Table 6**). Examination of RNA abundances of the lysosomal CLEAR network genes documented in HeLa cells^66^ revealed significant upregulation of *ACP5*, *ASAH1*, *CTSB*, *CTSD*, *CTSS*, *GAA*, *NPC2*, *PSAP* and *TPP1* (**Extended Data Fig. 5C, Table 7**). Next, we utilised gene set enrichment analysis (GSEA) to characterise up- and downregulated pathways in VPS35 KO cells (**Table 8**). This revealed enrichment of cellular component genes including endo-lysosomal genes, most notably subunits of the v-ATPase, as well as components of the mitochondrial respiratory chain complex; many of which are known TFEB target genes^67–70^ (**Extended Data Fig. 5D**). Moreover, there was a significant enrichment of genes associated with the KEGG pathways of Parkinson’s, Alzheimer’s, and Huntington’s diseases (**Extended Data Figs. 5E-G**).

**Figure 5.**
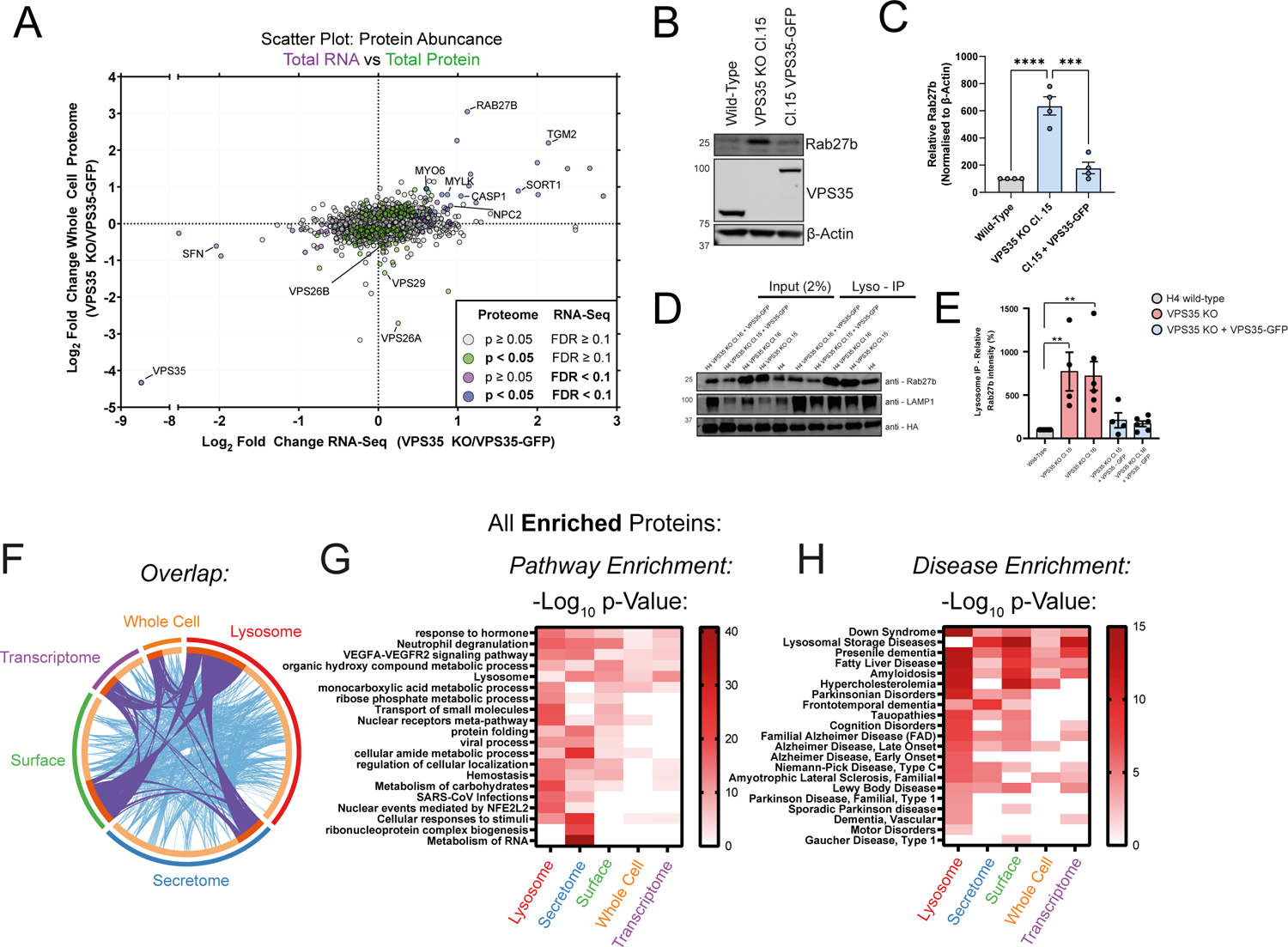
Correlative mapping of ‘omic’ datasets reveals transcriptional upregulation of Rab27b and enriched diseases. **(A)** Correlative analysis between RNA-Seq and the whole cell proteome reveals upregulation of a specific cohort of proteins. Scatter plot of VPS35 KO/VPS35-GFP RNA-Seq transcript abundances (x-axis) versus the protein abundances in the whole cell proteome (y-axis). Datapoints are coloured based on p-value scores in each experiment as denoted in the key. **(B-E)** Western blot and quantification of Rab27b protein levels in the whole cell lysates (B, C) and lysosomes (D, E) of wild-type, VPS35 KO, or VPS35-GFP rescue expression lines. For figure (C), n = 4 independent experiments, means ± SEM, ordinary one-way ANOVA comparison with Tukey’s multiple comparisons tests, wild-type vs VPS35 KO p = < 0.0001, VPS35 KO vs VPS35-GFP p = 0.0002, wild-type vs VPS35-GFP p = 0.4762. For (E), n = 5 independent experiments, means ± SEM, ordinary one-way ANOVA comparison with Dunnett’s multiple comparisons tests, * p < 0.05, ** p < 0.01, *** p < 0.001, **** p < 0.0001. **(F)** Circos plot of overlap of significantly enriched proteins in VPS35 KO samples compared to both wild-type and VPS35-GFP-expressing rescue cells across all datasets. **(G-H)** Pathway enrichment meta-analysis of significantly enriched categories (G) and associated DisGeneNET categories (H) in VPS35 KO cells compared to both wild-type and VPS35-GFP-expressing rescue cells across all datasets.

To refine our RNA-seq experiments, we performed whole-cell TMT proteomics of wild-type, VPS35 KO and VPS35-GFP rescue H4 samples (**Extended Data Fig. 6A, Tables 1-2)**. We correlated RNA transcript abundances with protein abundances to identify proteins that were upregulated at the transcriptional and proteomic level (**Fig. 5A**). Included within this cohort were *Rab27b*, *SORT1* and *NPC2*, which were also significantly perturbed in the VPS35 KO lysosomal proteome (**Figs. 3D-E**). The endosomal sorting of the lysosomal hydrolase receptor SORT1 is Retromer dependent^26, 71, 72^, and transcriptional upregulation may therefore reflect a compensatory mechanism to supply lysosomes with hydrolases in response to SORT1 mistrafficking and lysosomal dysfunction. Similarly, NPC2 upregulation may constitute a mechanism to restore cholesterol metabolism and egress from lysosomes due to perturb localisation of NPC2 (**Fig. 3H**).

**Figure 6.**
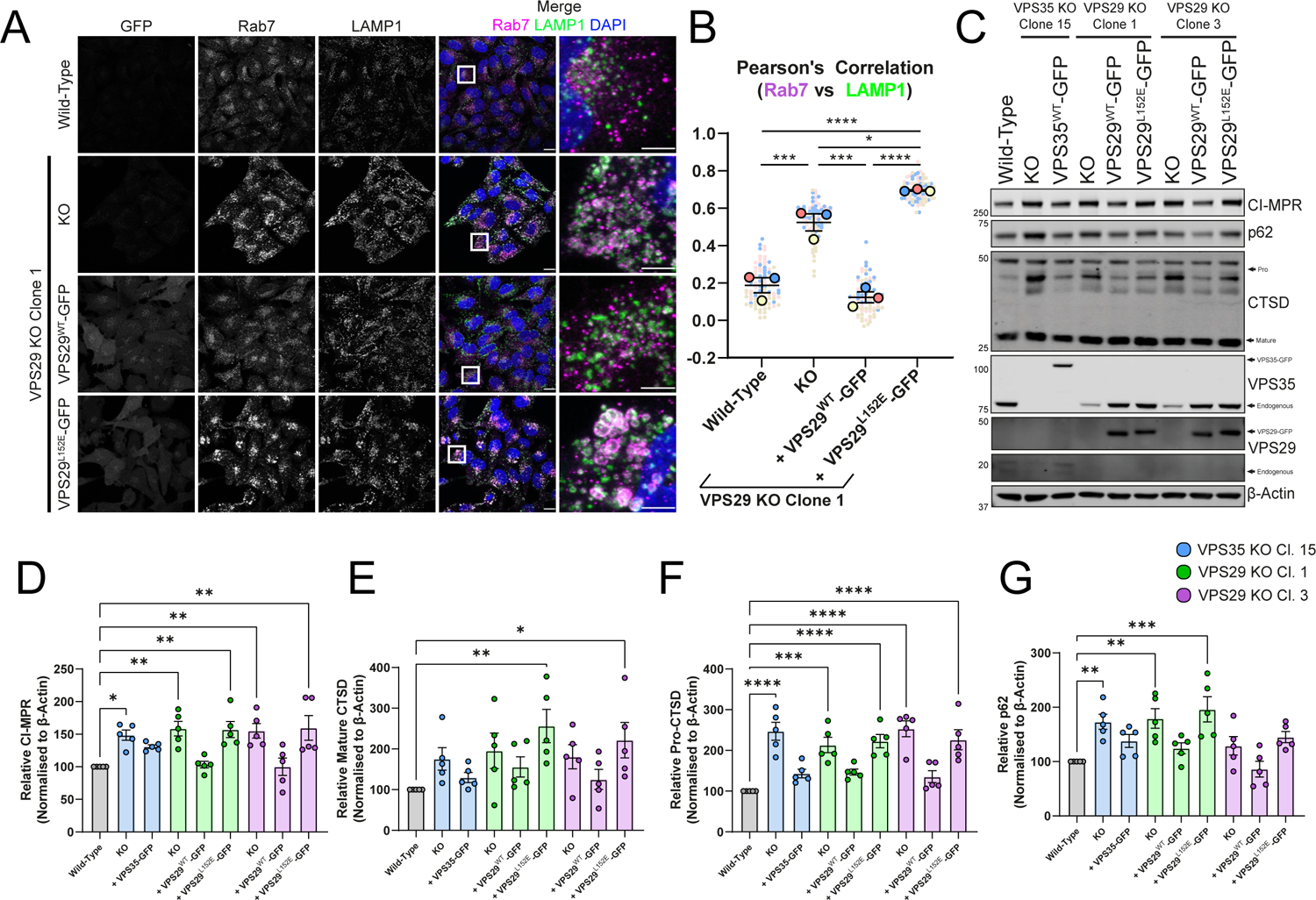
VPS29 KO H4 cells display a hyper-lysosomal recruitment of Rab7. **(A)** Deletion of VPS29 induces hyper-recruitment of Rab7 to lysosomes, which is rescued by re-expression VPS29^WT^-GFP but not VPS29^L215E^-GFP. Cell lines were fixed and immuno-stained for Rab7, LAMP1 and DAPI. Scale Bars: 20 µm and 5 µm in zoomed panels. **(B)** Rab7 labelling of lysosomes was quantified by measuring the Pearson’s correlation co-efficient between respective fluorescent signals over an n of 3 independent experiments. Means +/- SEM, one-way ANOVA with Tukey’s multiple comparisons tests. Wild-type vs VPS29 KO p = 0.0005, wild-type vs VPS29^WT^-GFP p = 0.5651, wild-type vs VPS29^L152E^-GFP p < 0.0001, VPS29 KO vs VPS29^WT^-GFP p = 0.0001, VPS29 KO vs VPS29^L152E^-GFP p = 0.308, VPS29^WT^-GFP vs VPS29^L152E^-GFP p < 0.0001. **(C-G)** Destabilisation of the Retromer trimer by VPS35 or VPS29 KO causes an increase in whole cell protein levels of CI-MPR, p62 and pro-CTSD. **(C)** Representative immuno-blot of endogenous protein levels derived from indicated cell lines. **(D-G)** Endogenous levels of indicated proteins were quantified relative to β-actin over n=5 independent experiments. Means +/- SEM, one-way ANOVA comparison with Dunnett’s multiple comparisons tests, * p < 0.05, ** p < 0.01, *** p < 0.001, **** p < 0.0001.

Rab27b was the most abundantly enriched protein in the VPS35 KO total cell proteome and was concomitantly enriched at the transcriptional level – this GTPase was also enriched in the lysosomal proteome (**Figs. 3D, 5A**). Rab27b regulates translocation of late endosomes to the cell periphery and their exocytosis^73^, and was recently shown to be transcriptionally regulated by the cytoprotective transcription factor Nrf2^74^. In Parkinson’s disease, higher expression of Rab27b has been reported in patient brain samples^75^, where it may promote the cell-to-cell transmission of pathogenic alpha-synuclein aggregates through a lysosomal exocytosis and re-uptake mechanism^75, 76^. We validated the increase and rescue of Rab27b in VPS35 KO cells by Western blotting of whole cell lysate and lysosome immunoprecipitates (**Figs. 5B-E**). Given the enrichment of lysosomal proteins, including APP, in the secretome of VPS35 KO cells, we speculate that the upregulation of Rab27b observed upon Retromer dysfunction may constitute a transcriptional link to stimulate lysosomal exocytosis.

Meta-analysis revealed disease signatures: including ‘Down Syndrome’, ‘Presenile Dementia’, ‘Amyloidosis’ and ‘Fatty Liver Disease’, which were unanimously enriched across all experimental approaches; and ‘Parkinsonian Disorders’, ‘Alzheimer Disease’, ‘Lewy Body Disease’, ‘Amyotrophic Lateral Sclerosis’ and ‘Niemann-Pick Disease’, among others, which were significantly enriched in multiple datasets (**Fig. 5F-H, Table 4**).

### Retromer Recruitment of Effector Proteins Govern Lysosomal Homeostasis

Finally, we returned to the LysoIP proteomics to seek insight into the mechanism(s) behind the swollen lysosomal phenotype. A possible origin for the multivariate phenotype observed in our datasets is the dysregulation of Rab7 due to impaired recruitment of the Retromer effector and Rab7 GAP TBC1D5^3–5^. Indeed, we observed a hyper-recruitment of Rab7 to LAMP1-positive membranes in VPS35 KO cells, in agreement with observations in HeLa cells from the Steinberg lab^3, 4^ (**Extended Data Fig. 7A**). TBC1D5 is recruited to Retromer though binding to VPS29^77^, which is impaired in the VPS29 L152E mutant^4^. We generated VPS29 KO H4 cells, which exhibited a comparable endo-lysosomal swelling and Rab7 hyper-recruitment to VPS35 KO cells, a phenotype rescued by re-expression of VPS29^WT^-GFP but not VPS29^L152E^-GFP (**Figs. 6A-C**). VPS35-GFP and VPS29-GFP rescued the accumulation of CI-MPR, CTSD and p62 observed in KO cells, but VPS29^L152E^-GFP failed to do so (**Fig. 6D-G**). Importantly, VPS35 protein levels were rescued in VPS29 KOs with VPS29^L152E^-GFP expression, indicating that Retromer assembly was unperturbed, thereby highlighting the importance of TBC1D5 recruitment mediated through VPS29 (**Extended Data Fig. 7B).** The binding of other VPS29 effector proteins, such as the VAMP7-interacting protein VARP, are also affected by the VPS29 L152E mutation^78^. We therefore cannot exclude additional consequences of this mutation, such as perturbed VAMP7-mediated fusion dynamics, contributing to lysosomal dysfunction. Taken together, these data establish a broad endo-lysosomal phenotype in Retromer-depleted H4 cells that arise, in part, from dysregulation of Rab7 nucleotide cycling.

**Figure 7.**
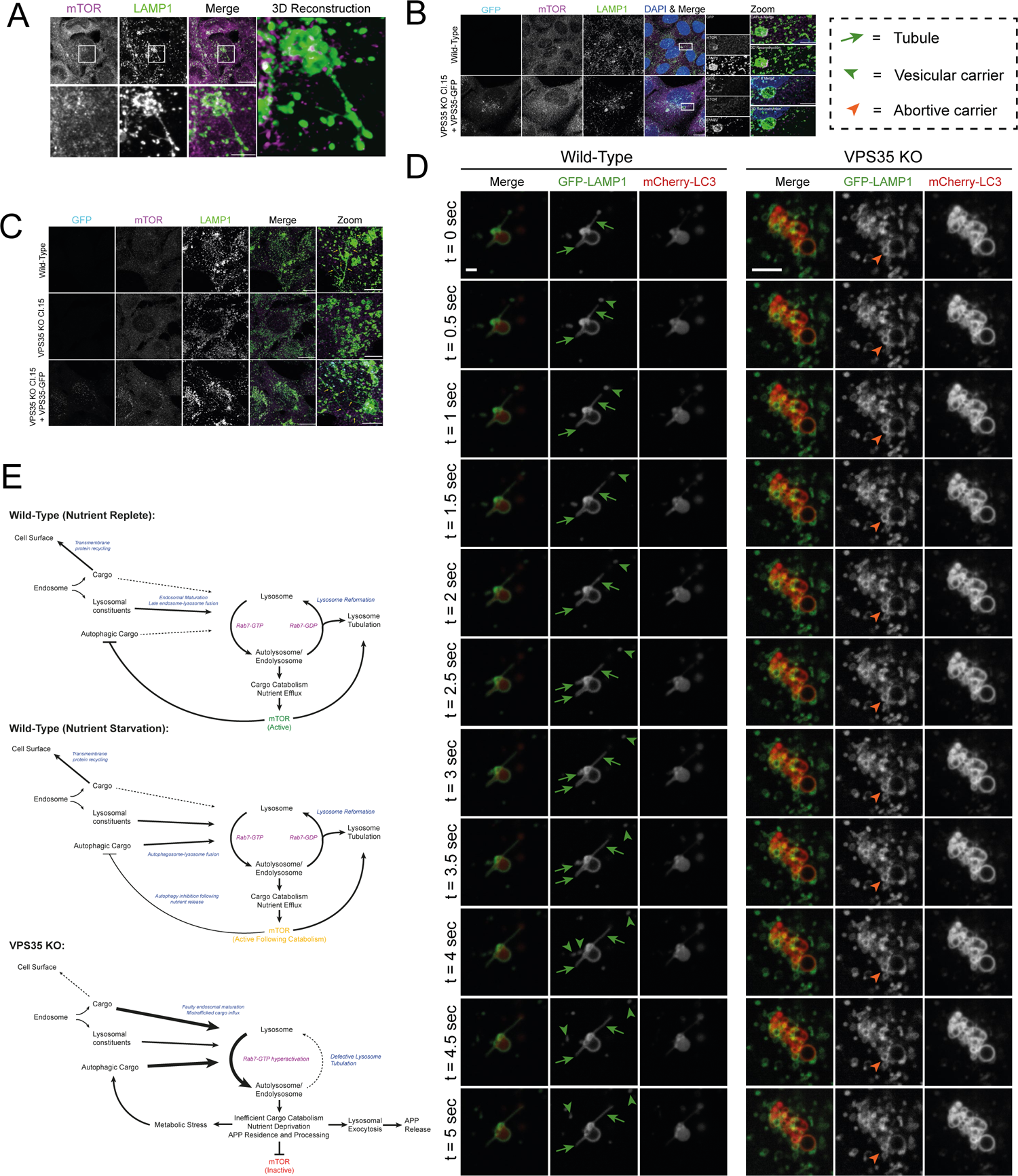
Perturbed autolysosome reformation induces defects to endo-lysosomal morphology in VPS35 KO. **(A)** Tubular resolution of H4 autolysosomes in response to amino acid starvation. H4 cells were depleted of amino acids for 2h prior to fixation and immuno-staining for mTOR and LAMP1. Magnified and 3D reconstructed panels depict mTOR and LAMP1 positive tubules emanating from autolysosomes. Scale Bars: 20 µm and 5 µm in zoomed panels. **(B)** Starvation induced autolysosome reformation in H4 is recapitulated in VPS35-GFP cells. H4 and VPS35-GFP cells were depleted of amino acids for 2h prior to fixation and immuno-staining for DAPI, mTOR and LAMP1. Magnified panels depict mTOR and LAMP1 decorated tubules emanating from autolysosomes co-labelled for VPS35-GFP. Scale Bars: 20 µm and 5 µm in zoomed panels. **(C)** Defective autolysosome resolution in VPS35 KO H4 cells. Cells were depleted of amino acids for 2 hours and refed in full media for 15 min prior to fixation and immuno-staining for mTOR and LAMP1. Zoomed panels show mTOR and LAMP1 tubules in H4 and VPS35 GFP compared to the LAMP1 positive, mTOR negative vesicular structures observed in VPS35 KO. Scale Bars: 20 µm and 5 µm in zoomed panels. **(D)** Representative time courses from live cell imaging of LAMP1 positive tubule formation and scission comparing events observed in wild-type and VPS35 KO cells. **(E)** Schematic depicting Rab7/mTOR-dependent activation of lysosome reformation under nutrient replete and starved conditions in wild-type cells. In VPS35 KO, reduced cargo recycling to the cell surface and increased flux into lysosomes, Rab7 hyperactivation, mTOR inactivation and perturbed lysosome tubulation illicit perturbed morphology and function of the endolysosomal network.

### VPS35 KO Cells Fail to Efficiently Undergo Lysosomal Reformation Events

While the hyper-recruitment and activation of Rab7 likely promotes the rate of late endosome and autophagosome fusion with lysosome, the decreased lysosomal enrichment of mTORC1, BORC, and PI(4)P metabolising enzymes is diagnostic for a defect in autophagic lysosomal reformation (ALR), a membrane tubulation events that resolve lysosomes back to their original size and density^79–81^. Amino acid starvation of wild-type and VPS35-GFP-expressing H4 cells induced a rapid dissociation of mTOR from lysosomes, which was recovered upon re-feeding in amino acid-replete DMEM **(Extended Data Fig. 8**). In VPS35 KO H4 cells, mTOR was already dissociated from lysosomes prior to amino acid starvation and failed to relocalise to lysosomes upon re-feeding **(Extended Data Fig. 8**), as shown recently in HeLa cells^3^.

Following periods of amino acid starvation extending beyond 60 minutes, we observed examples of ALR, defined by mTOR recruitment onto LAMP1-positive compartments with extended tubular structures (**Fig. 7A**). This phenomenon was observed in wild-type and VPS35-GFP-expressing cells during starvation and after amino acid re-feeding (**Figs. 7B-C**) but was not observed in VPS35 KO cells (**Fig. 7C**). We expressed GFP-RFP-LC3, a dual-colour autophagic flux reporter that loses GFP fluorescence within acidified autolysosomes. Expression of this reporter revealed extensive tubular networks of branches emanating from LAMP1-positive autolysosomes in wild-type cells upon amino acid starvation or re-feeding **(Extended Data Fig. 9**). Similar tubular events were far rarer in VPS35 KO cells, indicative of faulty ALR in the absence of mTOR-dependent nutrient sensing.

Live cell imaging revealed dynamic auto-lysosomal tubulation and fission events in wild-type cells expressing LAMP1-GFP and mCherry-LC3, which effectively serve to maintain compartment volume (**Fig. 7D, Supplementary Videos 1-6)**. In VPS35 KO cells, auto-lysosome membrane tubulation events, defined by tubulation of a LAMP1-GFP- and mCherry-LC3-positive compartment, were occasionally observed in VPS35 KO cells, but these were less frequent and less likely to undergo productive scission (**Fig. 7D, Supplementary Videos 7-12)**. Taken together, our data therefore reveal a defect in ALR as a major contributing factor in the swollen lysosomal phenotype observed in VPS35 KO cells, which mechanistically stems from the decreased lysosomal association of mTORC1, BORC, and PI(4)P-metabolising enzymes as identified through our integrated proteomics approach.

## DISCUSSION

### Towards a Global Understanding of the Neuroprotective Role of Retromer

Retromer has been associated with neurodegenerative disease since observation of reduced expression in Alzheimer’s disease patients and the identification of familial causative Parkinson’s disease mutations within the complex^11–16^. Our ‘omics-based approach has provided an unbiased analysis to highlight the detailed role of Retromer in safeguarding the functionality of the cell surface and maintaining lysosomal health. The deletion of this crucial sorting complex results in widespread dysfunction typified by loss of organelle identity, perturbed integral membrane protein sorting and transport, and inefficient lysosomal catabolism and lysosome resolution. With the emerging concept that small molecule compounds can be used to stabilise Retromer to enhance its neuroprotective function^82, 83^, our comprehensive cellular and molecular phenotyping of VPS34 KO H4 cells establishes a roadmap for future translational work to better understand the pathogenic link between Retromer-dependent regulation of lysosomal homeostasis and individual neurodegenerative diseases, and to inform diagnostic and therapeutic strategies.

### Dysfunctional Lysosomal Homeostasis in the Absence of Retromer

Lysosomal fusion and reformation dynamics are tightly regulated and essential for timely responsiveness to nutrient availability and catabolism. Following fusion of lysosomes with incoming endosomal or autophagic compartments, the resulting hybrid organelles (termed endo-lysosomes and auto-lysosomes, respectively) undergo a reformation process, whereby intraluminal contents are efficiently degraded, catabolites are exported out of the lysosome, and the resulting excess membrane and associated proteins are recycled through a tubulation process that is dependent on connectivity to the cytoskeleton. Through this mechanism, lysosomal size and number are controlled to facilitate cyclical rounds of fusion and reformation over the lifetime of a cell^84, 85^. Here, we demonstrate that besides its known role in autophagic flux^4, 6, 86^, Retromer is required for the ALR pathway to reform lysosomes following productive autophagic fusion events, a key step within the complex pathways that together maintain lysosomal homeostasis.

Following productive biogenesis of autophagosomes and lysosomal fusion, the delivered contents are efficiently catabolised, resulting in the efflux of nutrients. This is sensed by mTORC1, inducing a reactivation of mTORC1 activity to inhibit further autophagy and stimulate ALR^79^. Loss of mTOR association with lysosomes in VPS35 KO cells, and the mTORC1 depletion observed from LysoIP proteomics,suggests a perturbation to the lysosomal nutrient sensing system. Depletion of lysosomal solute transporters required for nutrient efflux, such as the L-glutamine transporter SLC38A7, may exacerbate this problem in catabolite transport and nutrient sensing at the lysosome (**Fig. 5J**).

mTORC1 signalling is required to recruit the BORC complex, a key regulator of lysosomal positioning and size^24, 87, 88^, and BORC is required for the generation of lysosomal tubulation during ALR^80^. Dynamic interconversion between PI(4)P and PI(4,5)P_2_ also contributes to the ALR tubulation process^81, 89–91^. Rab7-GTP hydrolysis is crucial for lysosomal reformation, since expression of constitutively active Rab7 or treatment with a non-hydrolysable GTP analogue inhibits ALR^79^. The decrease in PI4K2A, PIP4P1 and PIP4P2, and BORC components from VPS35 KO lysosomes, along with hyper-recruitment of active Rab7, provide the mechanistic insight into the observed decreased induction of ALR.

We therefore propose the following working model that underpins the lysosomal swelling phenotype observed in Retromer dysfunctional cells (**Fig. 7E**). Increased autophagic cargo influx and endosomal cargo missorting from the cell surface, combined with dysregulated endosomal maturation leads to inefficient degradation of lysosomal cargoes. Intraluminal contents accumulate within this environment, including APP. The impaired catabolism of these cargoes prevents effective nutrient efflux into the cytosol, which is required to initiate mTORC1- and BORC-dependent lysosome reformation events. Iterative rounds of fusion events and cargo delivery without effective resolution ultimately compound these problems, leading to the striking enlargement of hybrid organelles seen by light and electron microscopy. Endo- and auto-lysosomal exocytosis may constitute a cytoprotective response that releases undegraded contents from the cell, which in turn may facilitate cell-to-cell spreading of toxic contents such as pathogenic protein aggregates, which is emerging as a defining feature of neurodegenerative diseases^92^. Given that this broad lysosomal phenotype is observed in many models of both Retromer-dependent and Retromer-independent neurodegeneration, our data provide new insights into the neuroprotective role of Retromer in safeguarding lysosomal health.

### Lysosomal Exocytosis as a Compensatory Mechanism in VPS35 KO H4 Cells

The enrichment of soluble and transmembrane lysosomal proteins within the VPS35 KO ‘secretome’ is suggestive of lysosomal exocytosis. Retromer suppression has been associated with increased release of Aβ through exosomes in cell culture models^9^. Mass spectrometry of cerebrospinal fluid also recently revealed increased abundance of Tau in VPS35 KO mice and Alzheimer’s disease patient samples compared to controls^10^. In a recent study, Vps35 depleted *Drosophila* demonstrated APP accumulation in presynaptic neurons of the neuromuscular junction, and increased APP levels in extracellular postsynaptic vesicles^65^.

Lysosomal exocytosis has been proposed as a compensatory mechanism in response to lysosomal dysfunction^92^. For example, chemical perturbation of lysosomal homeostasis with ammonium chloride or bafilomycin A1 in SH-SY5Y cells leads to increased α-synuclein exocytosis and paracrine transfer to neighbouring cells^58^. This pathway may be particularly beneficial to neurons, whereby extracellular release of undegraded lysosomal contents alleviates lysosomal stress, and the released material can be internalised and degraded by neighbouring microglial cells in the brain. While this mechanism may be neuroprotective in the short-term, continued extracellular release of lysosomal material over the lifetime of an organism may contribute to the propagation and extracellular deposition and spread of pathogenic aggregates in later life as the ability of microglia to degrade this material diminishes. The discovery of Rab27b as one of the most abundantly enriched hits in VPS35 KO H4 cells, both at the transcript and protein level, provides a potential insight into how this process may be upregulated upon Retromer suppression.

Overall, our data emphasises the central importance of Retromer in controlling endolysosomal pathway function beyond its classical role in mediating the sequence-dependent retrieval of integral membrane proteins. This role of Retromer at the nexus of endolysosomal biology likely lies at the heart of its neuroprotective role and dysregulation in a number of neurodegenerative diseases. More generally, our integrated multiomic approach illustrates a powerful quantitative methodology through which to explore additional avenues for examining the dysregulation of the endo-lysosomal system observed in neurodegenerative disease.

## MATERIALS & METHODS

### Antibodies

#### Primary antibodies include

β-Actin (Sigma-Aldrich; A1978; clone AC-15; 1:2000 Western blot (WB)), APP (Abcam; Y188; ab32136; 1:200 immunofluorescence (IF)), ATP7A (Santa Cruz; D-9, sc-376467; 1:1000 WB), Cathepsin D (Proteintech; Clone, 21327-1-AP, 1:1000 WB), Cathepsin D (Merck; 219361; 1:200 IF), CI-MPR (Abcam; ab124767; clone EPR6599, 1:1000 WB, 1:400 immunofluorescence (IF)), EEA1 (Cell Signalling; 610456; clone 14; 1:200 IF), EGFR (Cell Signalling Technologies; 2232S; WB 1:1000), EGFR pY1068 (Cell Signalling Technologies; 3777S; WB1:1000), GFP (Roche; 11814460001; clones 7.1/13.1; 1:1000 WB), GLUT1 (Abcam; EPR3915; ab115730; 1:1000 WB; 1:50 IF), HA (Biolegend; 901502; Clone 16B12; 1:1000 WB, 1:200 IF), KEAP1 (Proteintech; 10503-2-AP; 1:1000 WB, 1:200 IF), LAMP1 (Developmental Studies Hybridoma Bank; AB_2296838; clone H4A3; 1:400 IF) LAMP1 (Abcam; ab21470; 1:200 IF), LAMP1 (Cell Signalling Technologies; CS4H11; 1:1000 WB), p62 (SQSTM1) (BD Transduction Laboratories; 610832; 1:1000 WB, 1:200 IF), Pmp70 (Sigma-Aldrich; 70-18; SAB4200181; 1:1000 WB), Rab7 (Abcam; EPR7589; ab137029; 1:200 IF), Rab27b (Proteintech; 13412-1-AP, 1:500 WB), TGN46 (Bio-Rad; AHP500G; 1:400 IF), TfnR (Santa Cruz; H68.4; sc-65883; 1:1000 WB), VPS29 (Santa Cruz; D-1; sc-398874; 1:500 WB), VPS35 (Abcam; ab157220; clone EPR11501(B); 1:1000 WB).

#### Secondary antibodies

For Western blotting, 680nm and 800nm anti-mouse and anti-rabbit fluorescent secondary antibodies (Invitrogen - 1:20,000). For immunofluorescence, 488nm, 568nm and 647nm AlexaFluor-labelled anti-mouse, anti-rabbit and anti-sheep secondary antibodies (Invitrogen - 1:400). 0.5 µg/mL 4’, 6-diamidino-2-phenylindole dihydrochloride (DAPI; Sigma-Aldrich, D8417) was added to secondary antibody mixtures to label DNA.

### Cell Culture

HeLa and HEK293T cells were sourced from the American Type Culture Collection (ATCC). H4 neuroglioma cells were a gift from Dr Helen Scott and Professor James Uney (University of Bristol). Clonal VPS35 KO HeLa and H4 cell lines were generated with the gRNA sequence 5′-GTGGTGTGCAACATCCCTTG-3′ targeting exon 5 of *VPS35*, ^18, 93^. VPS29 KO cells were generated by transfecting cells with gRNA sequences targeting the sequences 5’-GGACATCAAGTTATTCCAT-3’ and 5’-GGCAAACTGTTGCACCGGTG-3’ within exons 2 and 3 of *VPS29*, respectively.

Cells were grown in Dulbecco’s Modified Eagle Medium (DMEM; Sigma-Aldrich), supplemented with 10% (vol/vol) fetal bovine serum (FBS) (Sigma-Aldrich) and penicillin/streptomycin (Gibco). Cells were transduced with HIV-1-based lentiviruses for stable expression (construct of interest in pXLG3/pLVX/pLJC5 ^22^ plasmid backbone, and pCMV-dR8.91 packing plasmid) pseudotyped with vesicular stomatitis virus (VSV)-G envelope plasmid (pMDG2). HEK293T cells were transfected with the constituent plasmids using polyethyleneimine (PEI) transfection, then lentiviral particles were harvested after 48 hours. H4 cells were seeded into a plate, then transduced with lentivirus following adherence. For pLVX- and pLJC5-expressing cells, 3 μg/mL puromycin dihydrochloride was used for selection.

For amino acid starvation experiments, cells were plated the day before starvation. The culture media (DMEM containing 10% FBS and amino acids) was removed, followed by three PBS washes, and replaced with DMEM lacking amino acids and growth serum for the indicated timepoints. For re-feeding, the starvation media was removed and replaced with DMEM containing amino acids but lacking FBS.

### LysoIP

All equipment was pre chilled and all experimentation was performed at 4°C. Cells were washed twice in ice-cold PBS and harvested by scraping into 5 ml of KBPS (136mM KCl, 10mM KH_2_PO_4_ – pH to 7.5 using KOH) containing freshly added 5mM TCEP (Thermo #77720). Cells were pelleted by centrifugation at 270 x g for 10 min, re-suspended in 1ml of lysis buffer KPBS + TCEP and protease/phosphatase inhibitors, prior to mechanical lysis by 6 passages through a 23G needle. Cell debris was pelleted by centrifugation at 700 x g for 10 min. An aliquot of the lysate was removed to represent the whole cell lysate (treated with Triton TX-100 to a final concentration of 1% and an additional centrifugation at 18400 x g for 10 min to remove insoluble debris). Lysate volumes were re-adjusted to 1ml using lysis buffer and added to KPBS washed (x3) magnetic anti-HA beads (Thermo #88837) and gently rotated for 15 min. Beads were pelleted using a magnetic rack and three times washed in KPBS+TCEP for 5 mins with gentle rotation. Beads were pelleted on a magnetic rack and all trace of washing buffer removed, prior to re-suspension in RIPA buffer (10mM Triz pH7.5, 150mM NaCl, 1% TX100, 1% Deoxycholate, Protease and Phosphatase inhibitors) and incubation for 15 min with gentle rotation. Beads were pelleted on a magnetic rack and the eluate (solubilised lysosomal material) removed for subsequent analyses.

### Surface Biotinylation

All buffers were pre-chilled to 4°C. Cells were washed twice in ice-cold PBS prior to immersion in ice-cold PBS (pH7.7) containing 200µg/ml biotin (Thermo Scientific #A39258) for 30 min with gentle agitation at 4 ^O^C. To remove excess biotin, cells were washed in 1x PBS followed by 1x in Quench buffer (50mM Tris, 100mM NaCl, final pH7.5) prior to a 10 min incubation in quench buffer with gentle agitation. Cells were lysed by scraping in PBS (2% TX100 and protease inhibitor tablets) prior to pelleting of insoluble debris by centrifugation (14k for 10 min). An aliquot of the subsequent cleared lysate was retained to represent the whole cell fraction and the remainder added to pre-washed (in lysis buffer) streptavidin beads (Streptavidin sepharose – Cytiva #17511301). Precipitation of biotinylated cell surface proteins proceeded for 30 min at 4°C, prior to 1x wash in PBS + 1% TX100, 1x wash in PBS + 1% TX100 and 1M NaCl and a final wash in PBS. Biotin precipitated beads were pelleted by centrifugation and all traces of wash buffer removed prior to subsequent analyses.

### Quantitative Western Blotting

Bicinchoninic acid (BCA) assay (Pierce, 23225) or 660 nm assay (Pierce, 22662) was used to determine protein concentration according to the manufacturer’s instructions. NuPAGE 4-12% gradient Bis-Tris precast gels (Life Technologies, NPO336) were used for SDS-PAGE, followed by transfer onto methanol-activated polyvinylidine fluoride (PVDF) membrane (Immobilon-FL membrane, pore size 0.45 μm; Millipore, IPFL00010). Membrane was blocked, then sequentially labelled with primary and secondary antibodies. Fluorescence detected by scanning with a LI-COR Odyssey scanner and Image Studio analysis software (LI-COR Biosciences).

### Immunofluorescence Microscopy and Analysis

HeLa and H4 cells were seeded onto 13 mm coverslips the day before fixation. DMEM was removed, followed by two washes with PBS, then cells were fixed in 4% paraformaldehyde (PFA) (Pierce, 28906) for 20 minutes at room temperature. To visualise lysosomal tubules, cells were fixed in 8% PFA in 2X microtubule stabilization buffer (60 mM PIPES pH 6.8, 10 mM EGTA, 2 mM MgCl_2_) added directly to the cell culture media at a 1:1 volume ratio on a 37°C heat block, then returned to the tissue culture incubator for 15 minutes. Cells were permeabilised in 0.1% (w/v) saponin (Sigma-Aldrich, 47036) for 5 minutes followed by blocking with 1% (w/v) BSA, 0.01% saponin in PBS for 15 minutes. Coverslips were stained with primary antibodies for 1 hour, followed by secondary antibodies for 30 minutes, then mounted onto glass microscope slides with Fluoromount-G (Invitrogen, 00-4958-02).

Confocal microscope images were taken on a Leica SP5-II confocal laser scanning microscope attached to a Leica DMI 6000 inverted epifluorescence microscope or a Leica SP8 confocal laser scanning microscope attached to a Leica DM l8 inverted epifluorescence microscope (Leica Microsystems), with a 63x UV oil immersion lens, numerical aperture 1.4 (Leica Microsystems, 506192). For the Leica SP8 microscope, ‘lightning’ adaptive image restoration was used to generate deconvolved representative images.

Colocalisation and fluorescence intensity analysis was performed using Volocity 6.3 software (PerkinElmer) with automatic Costes background thresholding ^94^. Immunofluorescence images were prepared in Volocity 6.3. Lysosomal positioning quantification was performed in ImageJ as described in ^95^. Electron microscopy figures were prepared in ImageJ.

### Live cell imaging

Live-cell imaging was performed at 37°C with cells incubated in starvation media (formulated according to the Gibco recipe for high-glucose DMEM, omitting amino acids / FCS prior to filtration through a 0.22μm filter) or DMEM supplemented with 10% FCS in a CO2 buffered chamber. Fluorescent cells were imaged live on a Olympus Ixplore - SoRa spinning disk confocal system attached to a Olympus IX83 inverted epifluorescence microscope and a Hamamatsu sCMOS camera. Rapid switching between excitation/emission wavelengths facilitated a capture rate of ∼2 frames per second.

### Electron Microscopy

H4 cells were seeded onto 13 mm Thermanox Coverslips (Thermo Scientific) the day before fixation. 10nm BSA-gold (VWR) was ultracentrifuged at 100,000 x g for 1 hour at 4°C, the supernatant discarded, then the pellet was resuspended in 5 mL of complete DMEM media. The cell culture media was replaced with 10nm BSA-gold-containing media and cells were incubated at 37°C for 4 hours. Cells were fixed in a 2% paraformaldehyde, 2.5% glutaraldehyde and 0.1M sodium cacodylate solution for 30 minutes. Cells were then stained using 1% osmium tetroxide, 1.5% potassium ferrocyanide for 1 hour before staining was enhanced by incubation with 1% tannic acid in 0.1M cacodylate buffer for 45 minutes. Cells were washed, dehydrated through an ethanol series, and infiltrated with Epoxy propane (CY212 Epoxy resin:propylene oxide) before being infiltrated with full CY212 Epoxy resin and subsequently embedded atop pre-baked Epoxy resin stubs. Epoxy was polymerised at 65 °C overnight before Thermanox coverslips were removed using a heat-block. 70nm sections were cut using a Diatome diamond knife mounted to an ultramicrotome and sections collected to Pioloform-coated copper slot grids. Ultrathin sections were stained with lead citrate. An FEI Tecnai transmission electron microscope at an operating voltage of 80kV was used to visualise samples, mounted with a Gatan digital camera.

### Proteomics

#### Experimental Design

All proteomic experiments were performed with isobaric tandem mass tagging followed by LC-MS/MS quantitative mass spectrometry. For Lyso IP, 7 independent wild-type cells, 3 independent VPS35 KO Clone 15 and Clone 15 VPS35-GFP rescue and 4 independent VPS35 KO Clone 16 and Clone 16 VPS35-GFP rescue samples were quantified, producing 7 independent repeats of the wild-type vs VPS35 KO vs VPS35-GFP. For surface biotinylation, 6 independent wild-type cells, 3 independent VPS35 KO Clone 15 and Clone 15 VPS35-GFP rescue and 3 independent VPS35 KO Clone 16 and Clone 16 VPS35-GFP rescue samples were quantified, producing 6 independent repeats of the wild-type vs VPS35 KO vs VPS35-GFP. For the whole cell and ‘secretome’ proteomics, 3 independent wild-type samples, 1 VPS35 KO Clone 9 and Clone 9 VPS35-GFP, 1 VPS35 KO Clone 15 and Clone 15 VPS35-GFP and 1 VPS35 KO Clone 16 and Clone 16 VPS35-GFP samples were quantified, producing 3 independent repeats of the wild-type vs VPS35 KO vs VPS35-GFP experimental approach.

Samples for whole cell lysate analysis of H4 cells, cells were grown to confluency in a 10 cm plate, then lysed with 1% TX-100 lysis buffer and quantified with a BCA assay. The concentrations and volumes were normalised to a 200 µL volume of 2 mg/mL protein for each sample. To prepare samples for growth media ‘secretome’ analysis, H4 cells were grown in a 6-well plate in DMEM media without FBS for 16 hours. The medium was removed and centrifuged at 300 x g for 10 minutes at 4°C, then the supernatant was transferred to a fresh microcentrifuge tube and centrifuged at 2,000 x g for a further 10 minutes 4°C. The corresponding cells in the 6-well plate were lysed and quantified with a BCA assay to normalise media volumes.

#### TMT Labelling and High pH reversed-phase chromatography

For the cell surface proteome analysis, samples on beads were reduced (10mM TCEP, 55°C for 1h), alkylated (18.75mM iodoacetamide, room temperature for 30min.) and then digested from the beads with trypsin (2.5µg trypsin; 37°C, overnight). Alternatively, 50ug of each sample (whole cell lysate analysis), Bead eluates (LysoIP analysis), or media samples following concentration to 100ul using Amicon Ultra 3kDa cut-off centrifugal filters (Merck Millipore Ltd.) (secretome analysis) were reduced, alkylated and digested with trypsin, as described above. Following tryptic digestion, the resulting peptides were labelled with Tandem Mass Tag (TMT) ten plex reagents according to the manufacturer’s protocol (Thermo Fisher Scientific, Loughborough, LE11 5RG, UK) and the labelled samples pooled.

The pooled sample was evaporated to dryness, resuspended in 5% formic acid and then desalted using a SepPak cartridge according to the manufacturer’s instructions (Waters, Milford, Massachusetts, USA). Eluate from the SepPak cartridge was again evaporated to dryness and resuspended in buffer A (20 mM ammonium hydroxide, pH 10) prior to fractionation by high pH reversed-phase chromatography using an Ultimate 3000 liquid chromatography system (Thermo Scientific). In brief, the sample was loaded onto an XBridge BEH C18 Column (130Å, 3.5 µm, 2.1 mm X 150 mm, Waters, UK) in buffer A and peptides eluted with an increasing gradient of buffer B (20 mM Ammonium Hydroxide in acetonitrile, pH 10) from 0-95% over 60 minutes. The resulting fractions (15 for the whole cell lysate analysis, or 5 for the secretome, LysoIP or cell surface proteome analyses) were evaporated to dryness and resuspended in 1% formic acid prior to analysis by nano-LC MSMS using an Orbitrap Fusion Tribrid mass spectrometer (Thermo Scientific).

#### Nano-LC Mass Spectrometry

High pH RP fractions were further fractionated using an Ultimate 3000 nano-LC system in line with an Orbitrap Fusion Tribrid mass spectrometer (Thermo Scientific). In brief, peptides in 1% (vol/vol) formic acid were injected onto an Acclaim PepMap C18 nano-trap column (Thermo Scientific). After washing with 0.5% (vol/vol) acetonitrile 0.1% (vol/vol) formic acid peptides were resolved on a 250 mm × 75 μm Acclaim PepMap C18 reverse phase analytical column (Thermo Scientific) over a 150 min organic gradient, using 7 gradient segments (1-6% solvent B over 1min., 6-15% B over 58min., 15-32%B over 58min., 32-40%B over 5min., 40-90%B over 1min., held at 90%B for 6min and then reduced to 1%B over 1min.) with a flow rate of 300 nl min^−1^. Solvent A was 0.1% formic acid and Solvent B was aqueous 80% acetonitrile in 0.1% formic acid. Peptides were ionized by nano-electrospray ionization at 2.0kV using a stainless-steel emitter with an internal diameter of 30 μm (Thermo Scientific) and a capillary temperature of 275°C.

All spectra were acquired using an Orbitrap Fusion Tribrid mass spectrometer controlled by Xcalibur 2.1 software (Thermo Scientific) and operated in data-dependent acquisition mode using an SPS-MS3 workflow. FTMS1 spectra were collected at a resolution of 120 000, with an automatic gain control (AGC) target of 200 000 and a max injection time of 50ms. Precursors were filtered with an intensity threshold of 5000, according to charge state (to include charge states 2-7) and with monoisotopic peak determination set to peptide. Previously interrogated precursors were excluded using a dynamic window (60s +/-10ppm). The MS2 precursors were isolated with a quadrupole isolation window of 1.2m/z. ITMS2 spectra were collected with an AGC target of 10 000, max injection time of 70ms and CID collision energy of 35%.

For FTMS3 analysis, the Orbitrap was operated at 50 000 resolution with an AGC target of 50 000 and a max injection time of 105ms. Precursors were fragmented by high energy collision dissociation (HCD) at a normalised collision energy of 60% to ensure maximal TMT reporter ion yield. Synchronous Precursor Selection (SPS) was enabled to include up to 10 MS2 fragment ions in the FTMS3 scan.

#### Data Analysis

The raw data files were processed and quantified using Proteome Discoverer software v2.1 (Thermo Scientific) and searched against the UniProt Human database (downloaded January 2022; 178486 sequences) using the SEQUEST HT algorithm. Peptide precursor mass tolerance was set at 10ppm, and MS/MS tolerance was set at 0.6Da. Search criteria included oxidation of methionine (+15.995Da), acetylation of the protein N-terminus (+42.011Da) and Methionine loss plus acetylation of the protein N-terminus (−89.03Da) as variable modifications and carbamidomethylation of cysteine (+57.021Da) and the addition of the TMT mass tag (+229.163Da) to peptide N-termini and lysine as fixed modifications. Searches were performed with full tryptic digestion and a maximum of 2 missed cleavages were allowed. The reverse database search option was enabled and all data was filtered to satisfy false discovery rate (FDR) of 5%.

### RNA-Seq

6 independent wild-type samples, 2 VPS35 KO Clone 9 and Clone 9 VPS35-GFP, 2 VPS35 KO Clone 15 and Clone 15 VPS35-GFP and 2 VPS35 KO Clone 16 and Clone 16 VPS35-GFP samples were quantified, producing 6 independent repeats of the wild-type vs VPS35 KO vs VPS35-GFP experimental approach. H4 cells were grown to confluence in a 6-well plate. Media was removed and cells were washed twice with ice cold PBS. Cells were lysed and RNA was purified using the RNeasy kit (Qiagen) according to the manufacturer’s instructions. RNA concentration was measured using a NanoDrop 1000 machine (Thermo Fisher). Concentrations of all samples were normalised to 50 ng/µL.

Total RNA was quantified using the Qubit 2.0 fluorimetric Assay (Thermo Fisher Scientific). Libraries were prepared from 125 ng of total RNA using the NEGEDIA Digital mRNAseq research grade sequencing service (Next Generation Diagnostics srl)^96^ which included library preparation, quality assessment and sequencing on a NovaSeq 6000 sequencing system using a single-end, 100 cycle strategy (Illumina Inc.).

The raw data were analyzed by Next Generation Diagnostics srl proprietary 3’DGE mRNA-seq pipeline (v1.0) which involves a cleaning step by quality filtering and trimming, alignment to the reference genome and counting by gene (https://sourceforge.net/projects/bbmap/) ^97, 98^. We filtered out all genes having < 1 cpm in less than n_min samples. Differential expression analysis was performed using edgeR ^99^.

Gene set enrichment analysis (GSEA) of RNA-Seq data was performed using the GSEA software (UC San Diego and Broad Institute) and MSigDB database of gene sets. Specifically, the cellular compartment gene ontology gene sets (c5.go.cc.v7.2.symbols.gmt) and Kyoto Encyclopaedia of Genes and Genomes (KEGG) pathway gene sets (c2.cp.kegg.v7.2.symbols.gmt) were used for analysis ^100, 101^. Gene set networks from GSEA were visualised using Cytoscape 3.3 software with the Enrichment Map plug-in ^102^.

### Statistics and Bioinformatic Analysis

Raw files from mass spectrometry were quantified using Proteome Discoverer software v2.1 (Thermo Fisher). Peptides were searched against the UniProt human proteome database using the SEQUEST algorithm. For normalisation of mass spectrometry data, protein abundances were normalised based on total peptide amount for each experiment. Where proteins were identified and quantified by an identical group of peptides as the master protein of their protein group, these are designated ‘candidate master proteins’. We then used the annotation metrics for candidate master proteins retrieved from Uniprot to select the best annotated protein which was then designated as master protein. This enables us to infer biological trends more effectively in the dataset without any loss in the quality of identification or quantification. The mass spectrometry data were searched against the human Uniprot database retrieved on 2021-01-14, and updated with additional annotation information on 2021-11-15. To assemble the integrated datasets presented in Supplementary Tables 1 and 2, proteins were compared between datasets firstly using master protein accessions, and secondly using candidate master proteins, to ensure the best possible comparison. RNA data was integrated into the proteomics data using the biomaRt package in R ^103^.

For statistical analysis of differential protein abundance between conditions, standard t-tests were used. Volcano plots were plotted either the VolcaNoseR webapp ^104^. Typically, thresholds of log_2_ fold change of ±0.26 (corresponding to a 1.2-fold enrichment or depletion), and a -log_10_ p-value of 1.3 (corresponding to 0.05) were set, although these thresholds were adjusted based on assessment of data distributions for various experiments. Scatter plots were constructed in GraphPad Prism (LaJolla, CA) software.

Gene ontology analysis was performed using Metascape ^105^ to represent pathway enrichment, DisGeneNET category enrichment, and protein-protein interaction (PPI) networks, and the PANTHER classification system ^106^ was used to represent cellular component enrichment. Raw gene ontology output data is provided in **Table 3**. The dotted line overlaid on pathway enrichment graph represents a p = 0.05 statistical cut-off. PPI networks were visualised using Cytoscape 3.3 software with the Enrichment Map plug-in ^102^.

All statistical analysis was performed on data from a minimum of 3 independent experimental repeats. GraphPad Prism 9 (La Jolla, CA) software was used for statistical analysis of Western blot and confocal microscopy data.

Graphs were prepared in GraphPad Prism 9 or VolcanoseR. Individual datapoints represent independent experimental repeats. Graphs are plotted representing the mean value ± the standard error of the mean (SEM) for each experimental condition. *n* represents the number of independent experimental repeats. In all graphs, * = p < 0.05, ** = p < 0.01, *** = p < 0.001, **** = p < 0.0001.

---

**Extended Data Figure 1.**
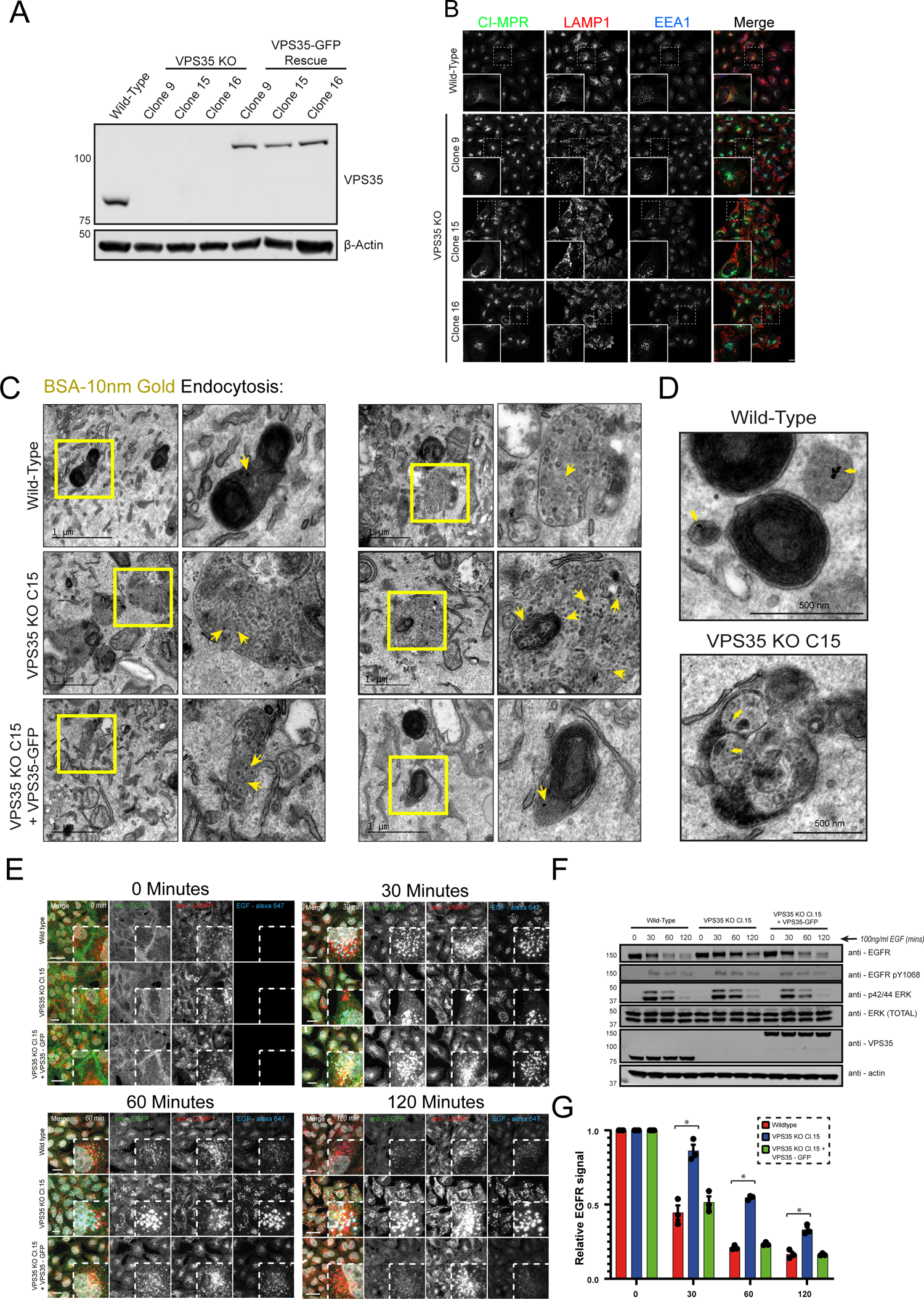
Validation of VPS35 KO Clones and Lysosomal Morphology Defects. **(A)** Western blot validation of VPS35-GFP expression in VPS35 KO H4 cells. The presence of the VPS35-GFP construct is indicated by increased molecular weight of VPS35, corresponding to GFP. **(B)** Immunofluorescence staining of three independent VPS35 KO clones with altered CI-MPR, LAMP1 and EEA1 compartment morphology. Scale bar = 20 µm, zoom scale bar = 5 µm. **(C-D)** Wild-type, VPS35 KO and VPS35-GFP-expressing rescue cells were incubated for 4 hours with 10nm BSA-gold prior to fixation and processing for electron microscopy. Transmission electron micrographs of internalized gold particles. Scale bars: 1 µm (C) and 500 nm (D). **(E)** EGFR is sorted to lysosomes but inefficiently degraded in VPS35 KO. Cells were serum starved to distribute EGFR at the plasma membrane prior to receptor activation via addition of EGF-Alexa-Fluor 647 (100 ng/ml) for the denoted time points and subsequent fixation and immuno-staining for EGFR and LAMP1. Scale bars: 20 µm. **(F)** The kinetics of EGFR degradation are perturbed in VPS35 KO relative to control and rescue cell lines. Cells were starved as above prior to stimulation with EGF (100 ng/ml) for the indicated time periods and immuno-blotting with anti-EGFR, EGFR pY1068, total ERK, phospho ERK (p42/44), VPS35 and β-actin. **(G)** Quantification of EGFR degradation time course over n=3 independent experiments. Means +/- SEM, one-way ANOVA with Tukey’s multiple comparisons tests. At 30 and 60 min EGF stimulation: wild-type vs VPS35 KO p = <0.0001 and VPS35-GFP vs VPS35 KO p = <0.0001, at 120 min: wild-type vs VPS35 KO p = 0.0037 and VPS35-GFP vs VPS35 KO p = 0.0002.

**Extended Data Figure 2.**
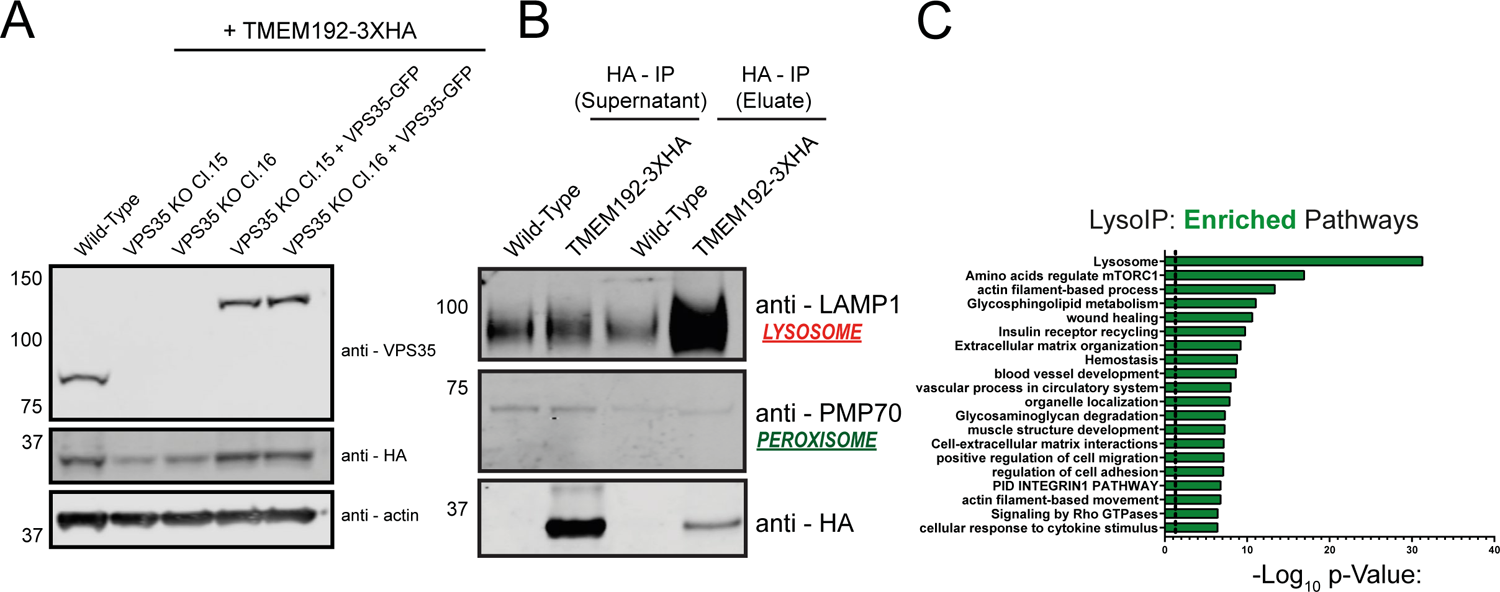
Development of LysoIP methodology. **(A)** Representative western blot depicting TMEM192-3xHA expression in cell lines. **(B)** LysoIP specifically enriches for lysosomal markers. Western blot probing for lysosomal and peroxisomal compartment markers in LysoIP samples derived from wild-type and TMEM192-3xHA expressing H4. **(C)** Pathway analysis of significantly enriched pathways in TMEM192-3xHA-expressing cells subjected to LysoIP relative to wild-type control H4 cells.

**Extended Data Figure 3.**
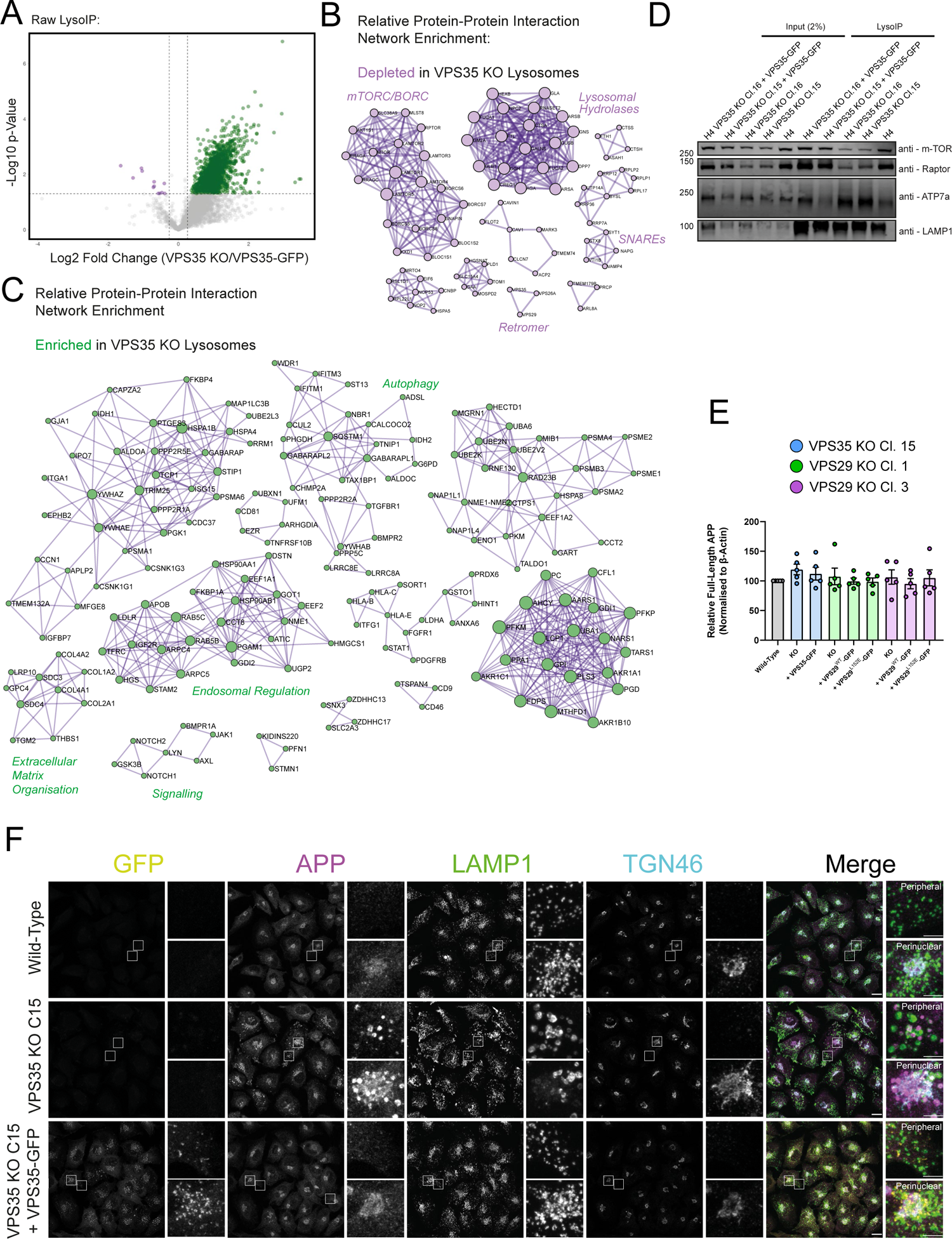
Additional LysoIP Proteomic Analysis. **(A)** Bulk lysosomal content is enriched in VPS35 KO cells. Volcano plot of raw VPS35 KO/VPS35-GFP protein abundances. 1024 and 6 proteins were significantly enriched and depleted, respectively, in VPS35 KO LysoIP compared to both wild-type and rescue samples (Log2 fold change ± 0.26, p < 0.05). **(B-C)** Protein-protein interaction networks of significantly depleted (B) and enriched (C) proteins in the normalised VPS35 KO LysoIP relative to wild-type and VPS35-GFP samples. **(D)** mTORC1 components dissociate from VPS35 KO lysosomes. Wild-type, VPS35 KO Clones 15 and 16 and corresponding VPS35-GFP rescues expressing TMEM192-3xHA were subjected to LysoIP followed by Western blotting. mTOR and Raptor levels decrease in VPS35 KO cells, whereas the retromer cell surface cargo ATP7a enriches in VPS35 KO lysosomes. **(E)** Quantification of full-length APP levels in wild-type, VPS35 KO and VPS29 KO cells with corresponding rescue construct expression, displayed in Figure 4K. n = 4 independent repeats. **(F)** APP accumulates within lysosomes in VPS35 KO cells. Immunofluorescence staining of wild-type, VPS35 KO Clone 15 and Clone 15 VPS35-GFP rescue cells stained with anti-APP-LAMP1 and -TGN46 antibodies. Insets depict examples of perinuclear APP (centred on the *trans*-Golgi network (TGN)) and peripheral APP. Scale bar = 20 µm, zoom scale = 5 µm.

**Extended Data Figure 4.**
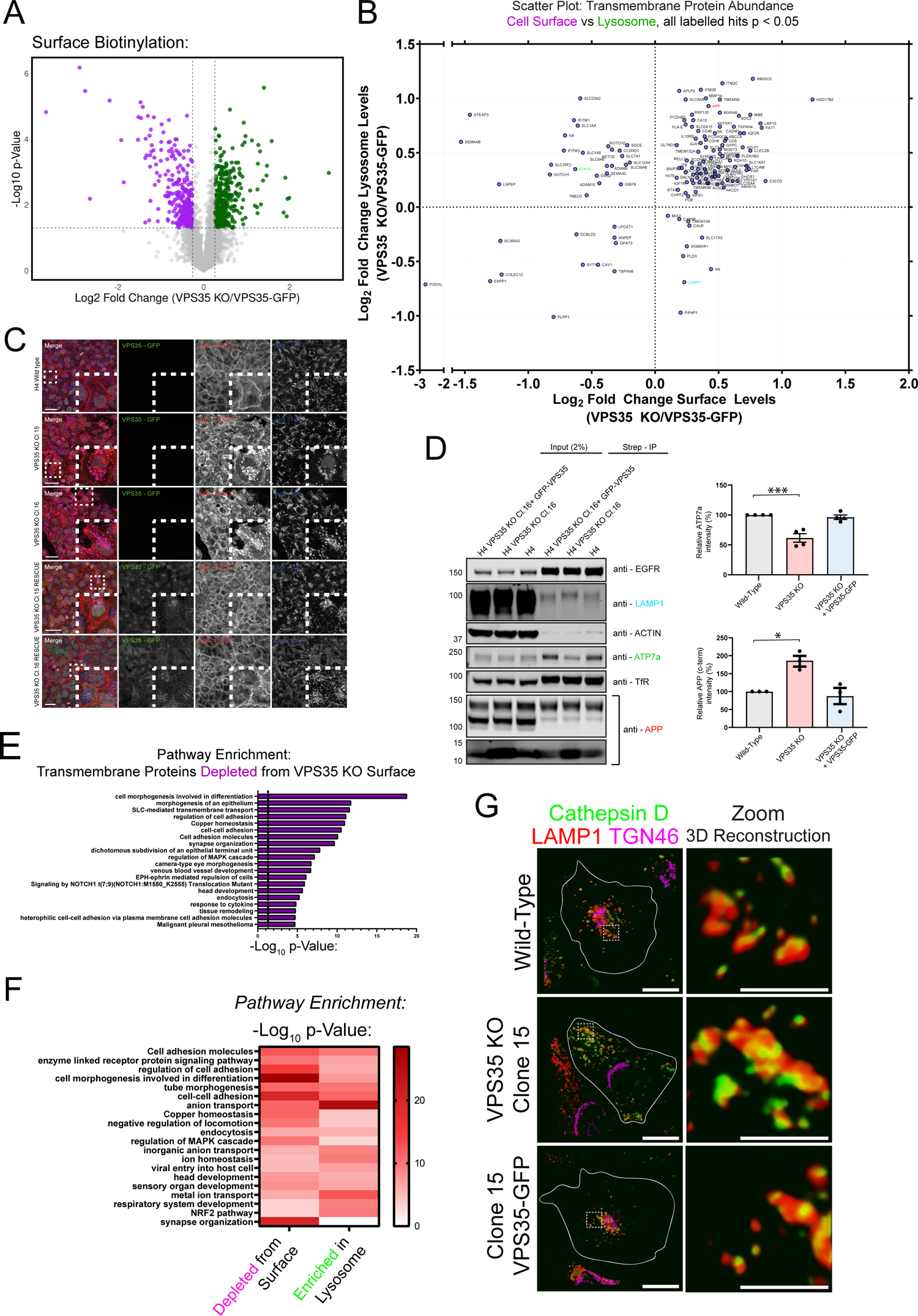
Proteomic Profiling of the VPS35 KO Cell Surface and Lysosomal Proteomes. **(A)** Volcano plot of VPS35 KO/VPS35-GFP protein abundances in the cell surface proteome. 270 and 199 proteins were significantly enriched and depleted, respectively (Log2 fold change ± 0.26, p < 0.05). **(B)** Scatter plot of VPS35 KO/VPS35-GFP protein abundances in the cell surface proteome (x-axis) versus the LysoIP proteome (y-axis). Only significantly proteins in both experiments are displayed and labelled (Log_2_ fold change ± 0.26, p < 0.05). **(C)** Imaging of endogenous GLUT1 steady-state distribution establishing that re-expression of VPS35-GFP rescues the missorting to lysosomes observed in VPS35 KO H4 cells. **(D)** Cell surface abundance of APP-CTF is elevated in VPS35 KO H4 cells. Cell surface precipitates from indicated cell lines were immuno-blotted using anti – APP (full length and CTF), EGFR, TfR and LAMP1 (confirmation of enrichment of surface proteome), β-actin and ATP7a. Quantification of APP-CTF (n=3) and ATP7a (n=4) cell surface abundances (means +/- SEM, one-way ANOVA with Dunnett’s multiple comparisons tests, comparisons with wildtype, p = * p < 0.02, *** p < 0.0005). **(E)** Pathway analysis of significantly depleted pathways in the VPS35 KO transmembrane cell surface proteome relative to wild-type and VPS35-GFP rescue controls. **(F)** Pathway enrichment analysis of significantly depleted pathways represented by transmembrane proteins significantly depleted from the VPS35 KO cell surface proteome, and transmembrane proteins significantly enriched in the VPS35 KO LysoIP proteome relative to wild-type and VPS35-GFP rescues. **(G)** Cathepsin D localises to LAMP1-positive compartments in VPS35 KO H4 cells. Immunofluorescence microscopy of wild-type, VPS35 KO and VPS35-GFP rescue H4 cells stained with anti-CTSD, -LAMP1 and -TGN46 antibodies. Scale bar = 20 µm, zoom 3D reconstruction scale bar = 5 µm.

**Extended Data Figure 5.**
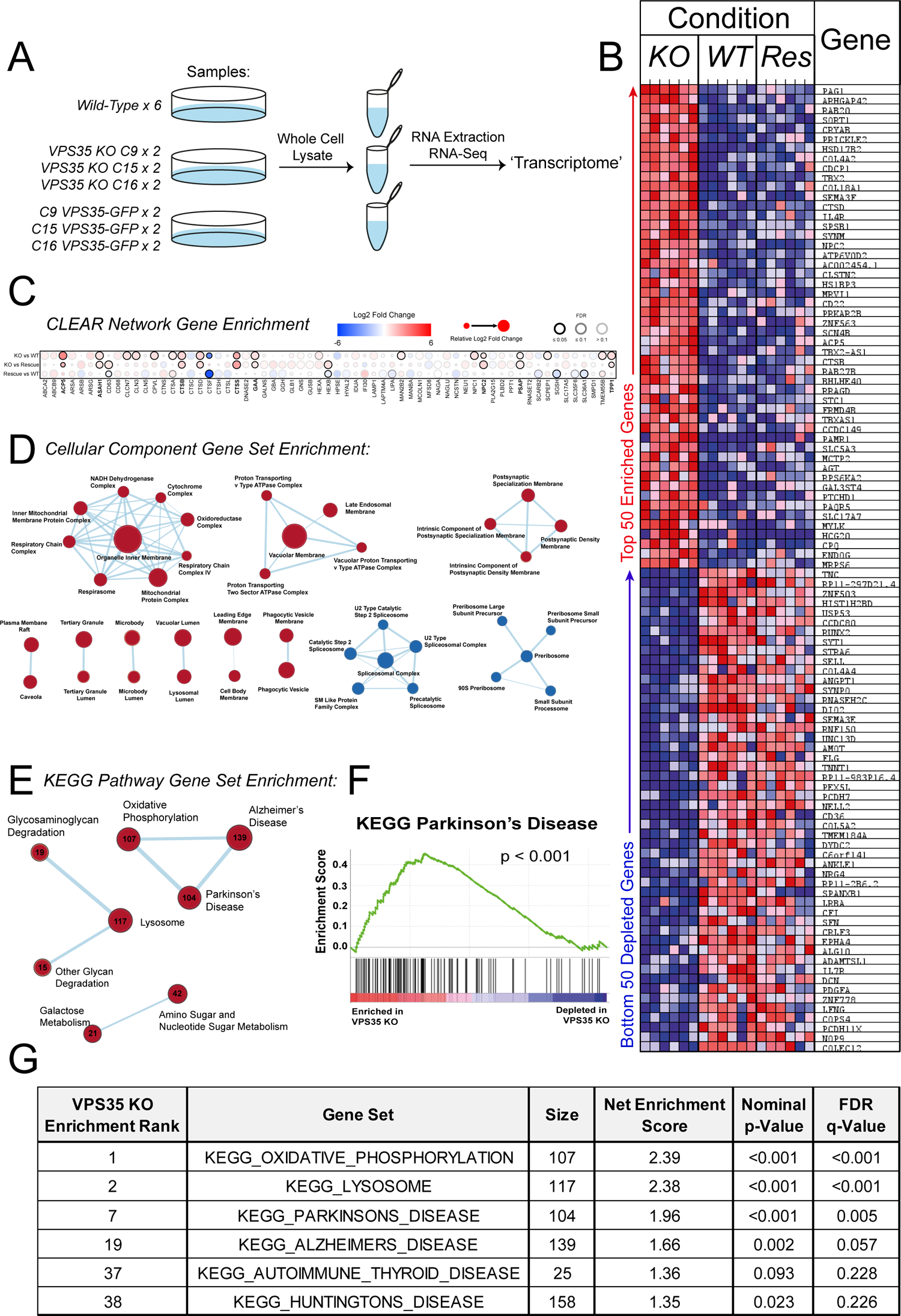
Transcriptional Profiling of VPS35 KO Cells. **(A)** Schematic of the RNA-seq experimental design. 6 wild-type samples, 6 VPS35 KO samples (2 each of VPS35 KO clones 9, 15 and 16) and 6 VPS35-GFP rescue samples (2 each of VPS35-GFP rescue clones 9, 15 and 16) were analysed. **(B)** Heatmap depicting the top 50 up- and down-regulated genes in VPS35 KO samples relative to both wild-type and rescue cells. **(C)** Dot-plot depicting the relative fold changes of CLEAR network gene transcripts in VPS35 KO cells relative to wild-type and VPS35-GFP rescue cells. Heatmap scale and FDR scores are indicated by colour of dots and rings, respectively. **(D)** Network analysis of significantly enriched cellular component gene sets. Red circles denote enriched categories, and blue circles denote depleted categories in VPS35 KO H4 cells. Circle size represents the number of enriched/depleted genes belonging to each gene set within the dataset. **(E)** Network analysis of significantly enriched KEGG pathway gene sets, presented as in (D). Circles are annotated with the number of enriched genes within each gene set. **(F)** Representative enrichment score plot genes enriched in the ‘Parkinson’s Disease’ KEGG pathway. **(G)** Table of selected significantly enriched KEGG pathway gene sets, depicting their rank, net enrichment score, size, and statistics. The full list of gene set enrichment is depicted in **Table 8**.

**Extended Data Figure 6.**
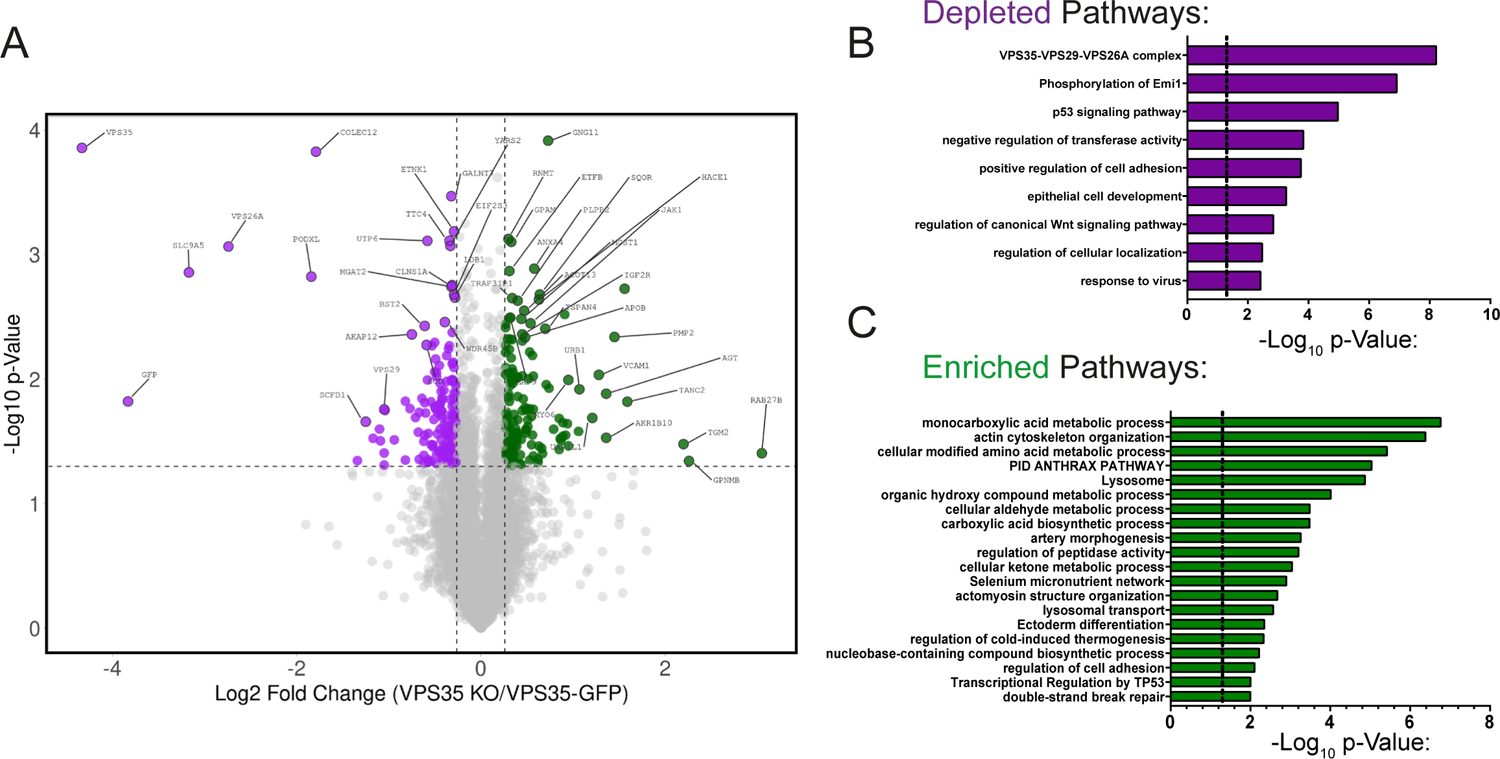
Supplementary Whole Cell Proteome and Meta-Analysis. (A) Volcano plot of VPS35 KO/VPS35-GFP protein abundances in the whole cell proteome. 71 and 59 proteins were significantly enriched and depleted, respectively (Log2 fold change ± 0.26, p < 0.05). **(B-C)** Pathway analysis of significantly depleted (B) or enriched (C) pathways in the VPS35 KO whole cell proteome.

**Extended Data Figure 7.**
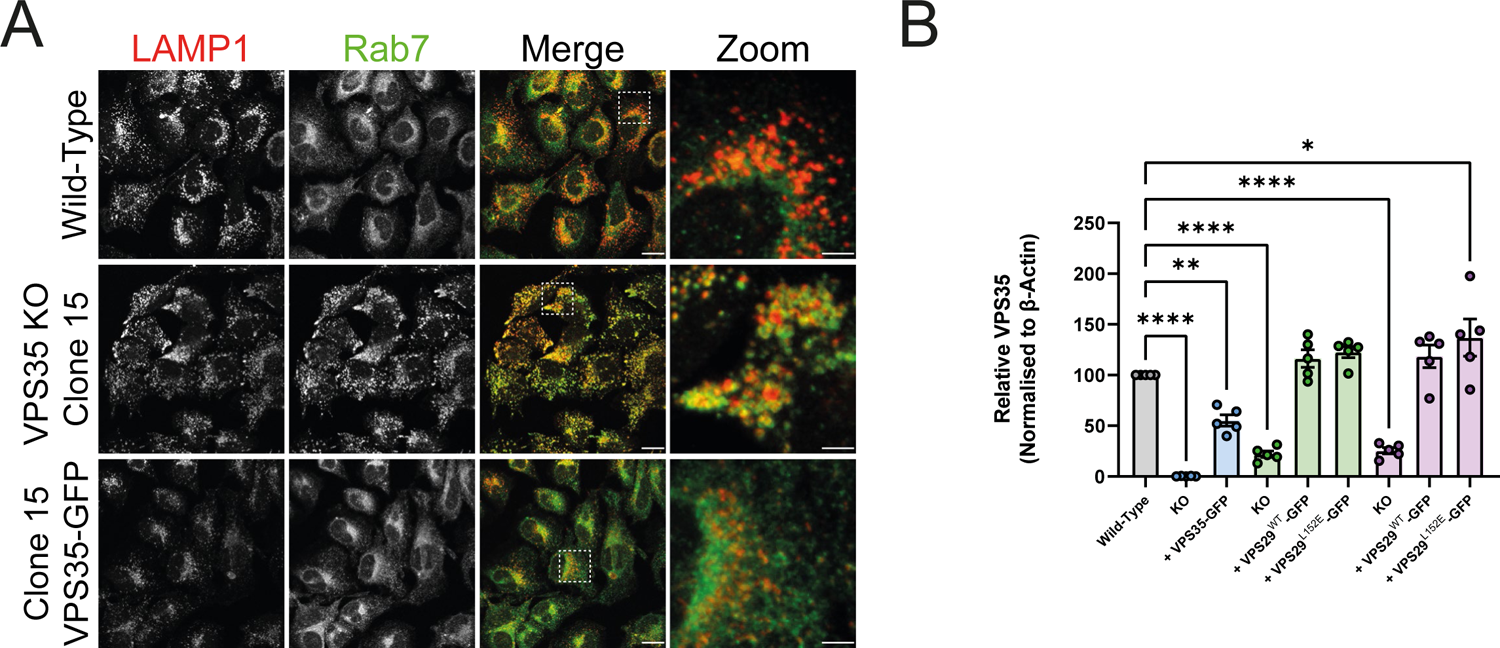
VPS35 KO H4 cells display hyper-recruitment of Rab7 to LAMP1-positive lysosomes. **(A)** VPS35 KO cells exhibit Rab7 hyper-recruitment to LAMP1-positive compartments. Immunofluorescence staining of wild-type, VPS35 KO Clone 15, and Clone15 VPS35-GFP-expressing cell stained with anti-Rab7 and -LAMP1 antibodies. Scale bar = 20 µm, zoom scale = 5 µm. **(B)** Quantification of VPS35 levels of indicated proteins were quantified relative to β-actin over n=5 independent experiments, displayed in Figure 2F. Means +/- SEM, one-way ANOVA comparison with Dunnett’s multiple comparisons tests, * p < 0.05, ** p < 0.01, *** p < 0.001, **** p < 0.0001.

**Extended Data Figure 8.**
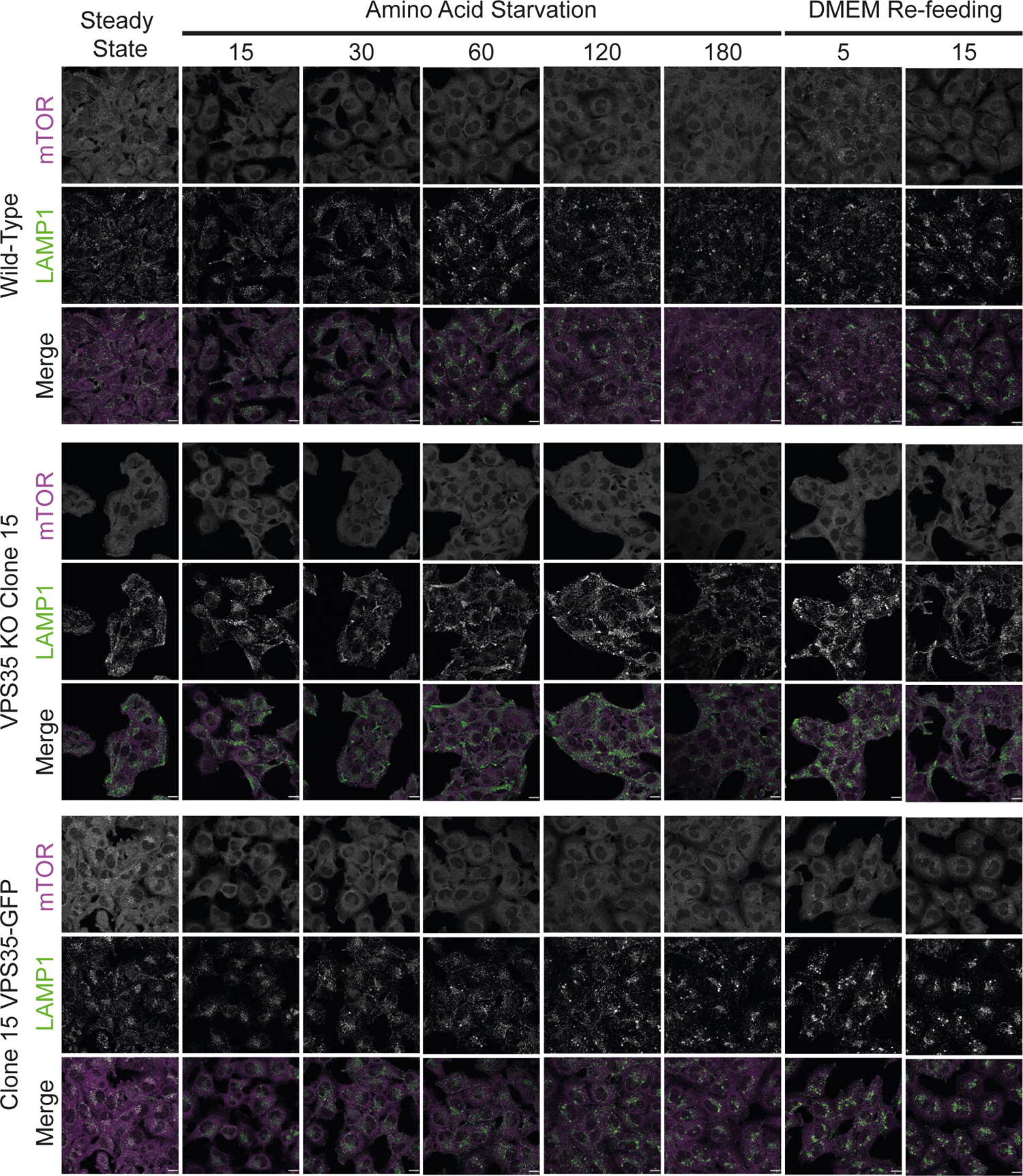
Impaired mTOR Recruitment in VPS35 KO H4 Cells Lysosomal association of mTOR is unresponsive to amino acid starvation and re-feeding in VPS35 KO. Cells were amino acid starved prior to re-feeding in DMEM for the indicated time periods, then fixed and immuno-stained for mTOR and LAMP1. Scale bars = 20 µm.

**Extended Data Figure 9.**
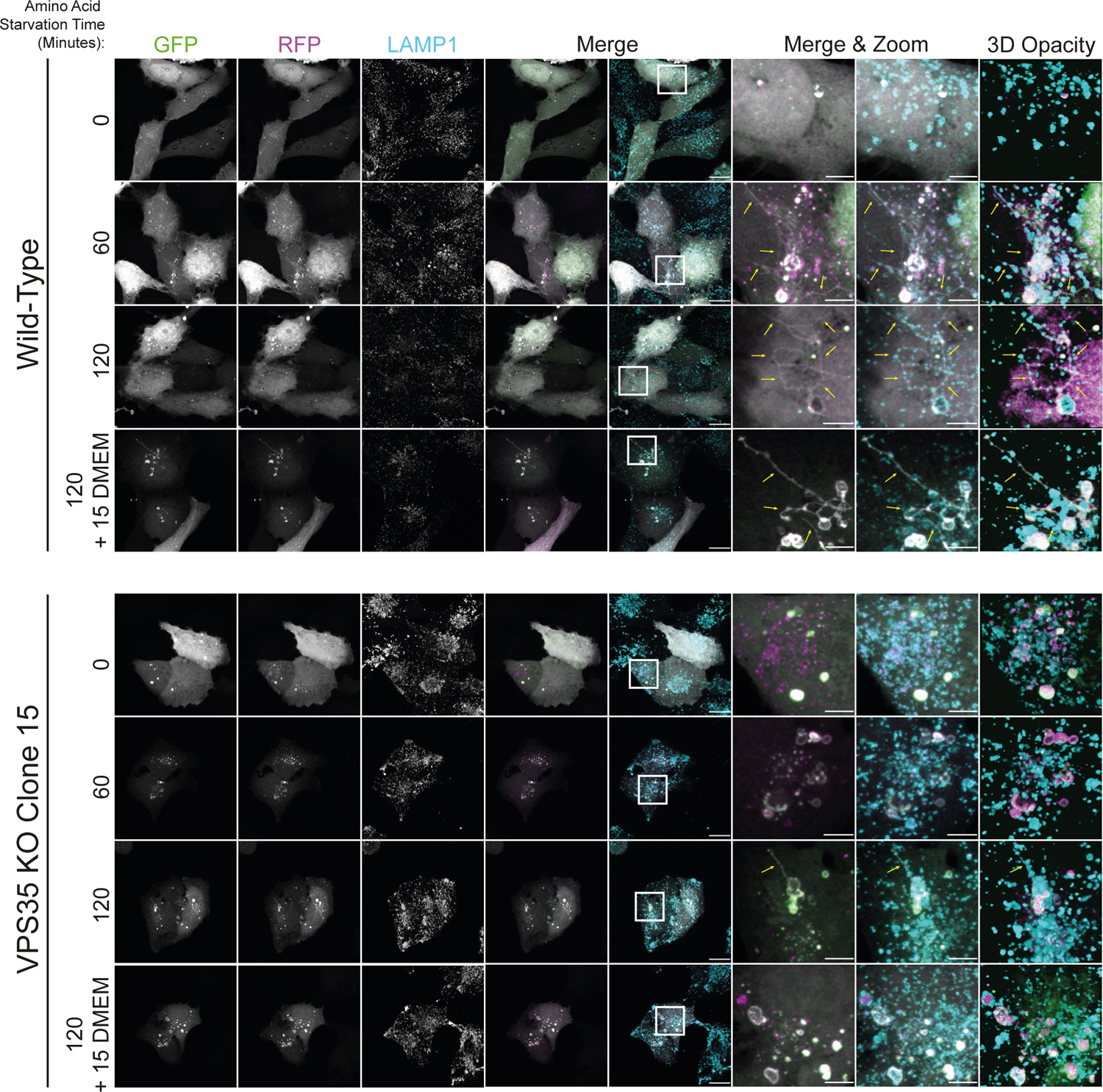
Defective Tubulation of Autophagic Lysosomes in VPS35 KO Cells LC3-positive autophagic lysosomes display abundant tubulation in following starvation or re-feeding in wild-type cells, whereas these events are less common in VPS35 KO H4 cells. Cells were transfected with a GFP-RFP-LC3 dual reporter construct and starved prior to re-feeding in DMEM for the indicated time periods, then fixed and immuno-stained for LAMP1. Scale bars = 20 µm, zoom scale = 5 µm.

## Table Legends

**Table 1:** Integrated Raw Proteomics and RNA-Seq Data from Wild-Type, VPS35 KO and VPS35-GFP-expressing H4 Neuroglioma Cells. Data were not normalised. Output data is displayed for each protein across all independent experiments, including gene ontology classifications, peptide counts, protein coverage, Log_2_ fold change, t-test p-values and false discovery rate (FDR). This dataset was used for analysis of raw abundances in LysoIP and growth media ‘secretome’ datasets.

**Table 2:** Integrated Normalised Proteomics and RNA-Seq Data from Wild-Type, VPS35 KO and VPS35-GFP-expressing H4 Neuroglioma Cells. Data were normalised based on total peptide amount for each experiment. Output data is displayed for each protein across all independent experiments, including gene ontology classifications, peptide counts, protein coverage, Log_2_ fold change, t-test p-values and false discovery rate FDR. This dataset was used for analysis of relative abundances in LysoIP, surface biotinylation, total cell proteome and RNA-Seq datasets.

**Table 3:** Metascape Pathway Enrichment Outputs from Proteomics Data. The top 20 enriched pathways for significantly enriched or depleted proteins across the datasets are provided, alongside p-value scores and the list of proteins within each category.

**Table 4:** Metascape DisGeneNET Enrichment Outputs from Proteomics Data. Outputs from 3 comparisons are displayed: proteins significantly enriched in VPS35 KO LysoIP proteomics dataset; significantly enriched proteins across all datasets; and significantly depleted proteins across all datasets. In all cases, the raw output with all diseases is displayed, and the selected outputs displayed on graphs in the manuscript are shown in a separate tab.

**Table 5:** Cellular Component Gene Ontology Analysis of the VPS35 KO ‘Secretome’. Significantly enriched proteins from VPS35 KO samples (Log_2_ fold change > 1, p < 0.05), were analysed using PANTHER gene ontology software. All cellular component gene ontology categories are shown, with their respective fold enrichments and p-value scores.

**Table 6:** RNA-Seq Quantification of Global Transcript Abundances. RNA transcripts are displayed along with their abundances in wild-type, VPS35 KO or VPS35-GFP-expressing H4 cells. Log_2_ fold changes and FDR values are displayed.

**Table 7:** RNA-Seq Quantification of CLEAR Network Genes. CLEAR network genes are displayed alongside their corresponding RNA-Seq Log_2_ fold changes and FDR values.

**Table 8:** Gene Set Enrichment Analysis (GSEA) of RNA-Seq Data. Cellular component and Kyoto Encyclopaedia of Genes and Genomes (KEGG) pathway gene sets significantly enriched in VPS35 KO RNA-Seq datasets are displayed with enrichment scores and statistics.

## Supplementary Video Legends

**Supplementary Videos 1-6 Autolysosome Tubulation in Wild-Type H4 Cells**

Spinning disk live fluorescence microscopy of a wild-type H4 cell co-transfected with LAMP1-GFP and mCherry-LC3 (Supplementary Movies 1-3) and a corresponding zoom region of an autolysosome tubulation events (Supplementary Movies 4-6). Supplementary Videos 1 and 4 show merged GFP and mCherry channels, Supplementary Videos 2 and 5 display GFP only, and Supplementary Videos 3 and 6 display mCherry only.

**Supplementary Videos 7-12 Perturbed Autolysosome Dynamics in VPS35 KO Cells**

Spinning disk live fluorescence microscopy of a VPS35 KO H4 cell co-transfected with LAMP1-GFP and mCherry-LC3 (Supplementary Movies 7-9) and a corresponding zoom region of a cluster of autolysosome compartments (Supplementary Movies 10-12). Supplementary Videos 7 and 10 show merged GFP and mCherry channels, Supplementary Videos 8 and 11 display GFP only, and Supplementary Videos 9 and 12 display mCherry only.

## Acknowledgements

We thank the Wolfson Bioimaging Facility and Bristol Proteomics Facility for their support. px458 plasmids targeting exons 2 and 3 of *VPS29* were cloned by Dr Kerrie McNally. P.J.C. was supported by the Wellcome Trust (104568/Z/14/Z and 220260/Z/20/Z), the Medical Research Council (MRC) (MR/L007363/1 and MR/P018807/1), the Lister Institute of Preventive Medicine, and the award of a Royal Society Noreen Murray Research Professorship (RSRP/R1/211004). J.L.D was supported by a Wellcome Trust Dynamic Molecular Cell Biology PhD Studentship (203959/Z/16/Z). J.R.E is supported by a Sir Henry Dale Fellowship jointly funded by the Wellcome Trust and the Royal Society (216370/Z/19/Z). This work was supported by Italian Telethon Foundation (TIGEM institutional grant), European Research Council H2020 AdG (LYSOSOMICS 694282 to A.B.) and Associazione Italiana per la Ricerca sul Cancro A.I.R.C. (IG-22103 to A.B.).

## Disclosure of Potential Conflict of Interest

A.B. is cofounder of CASMA Therapeutics and Advisory board member of Avilar Therapeutics and Coave Therapeutics. D.C and A.B are founders, shareholders, and consultants of Next Generation Diagnostic srl. S.R and L.D.F are employees of Next Generation Diagnostic srl.

## Contributions

J.L.D, C.M.D and P.J.C conceived the study. J.L.D, C.M.D, S.R, L.D.F, D.C, J.R.E and K.J.H performed experiments. J.L.D, C.M.D, P.A.L, S.R, L.D.F, D.C, S.J.C, and K.J.H analysed data. A.B, J.R.E and P.J.C acquired funding. P.J.C supervised work. J.L.D, C.M.D, and P.J.C wrote the initial manuscript with all authors editing and approving the final text.

## References

1. Cullen, P. J. & Steinberg, F. To degrade or not to degrade: mechanisms and significance of endocytic recycling. Nat. Rev. Mol. Cell Biol. 19, 679–696 (2018).

2. Seaman, M. N. J., McCaffery, J. M. & Emr, S. D. A membrane coat complex essential for endosome-to-Golgi retrograde transport in yeast. J. Cell Biol. 142, 665–681 (1998).

3. Kvainickas, A. et al. Retromer and TBC1D5 maintain late endosomal RAB7 domains to enable amino acid–induced mTORC1 signaling. J. Cell Biol. 218, 3019–3038 (2019).

4. Jimenez-Orgaz, A. et al. Control of RAB7 activity and localization through the retromer-TBC1D5 complex enables RAB7-dependent mitophagy. EMBO J. e201797128 (2017) doi:10.15252/embj.201797128.

5. Seaman, M. N. J. J., Harbour, M. E., Tattersall, D., Read, E. & Bright, N. Membrane recruitment of the cargo-selective retromer subcomplex is catalysed by the small GTPase Rab7 and inhibited by the Rab-GAP TBC1D5. J. Cell Sci. 122, 2371–2382 (2009).

6. Carosi, J. M. et al. Retromer regulates the lysosomal clearance of MAPT/tau. Autophagy 00, 1–21 (2020).

7. Bi, F., Li, F., Huang, C. & Zhou, H. Pathogenic mutation in VPS35 impairs its protection against MPP(+) cytotoxicity. Int. J. Biol. Sci. 9, 149–55 (2013).

8. Tang, F. L. et al. VPS35 in dopamine neurons is required for endosome-to-golgi retrieval of Lamp2a, a receptor of chaperone-mediated autophagy that is critical for α-synuclein degradation and prevention of pathogenesis of Parkinson’s disease. J. Neurosci. 35, 10613–10628 (2015).

9. Sullivan, C. P. et al. Retromer disruption promotes amyloidogenic APP processing. Neurobiol. Dis. 43, 338–345 (2011).

10. Simoes, S. et al. Tau and other proteins found in Alzheimer’s disease spinal fluid are linked to retromer-mediated endosomal traffic in mice and humans. Sci. Transl. Med. 12, (2020).

11. Small, S. A. et al. Model-guided microarray implicates the retromer complex in Alzheimer’s disease. Ann. Neurol. 58, 909–919 (2005).

12. Zimprich, A. et al. A mutation in VPS35, encoding a subunit of the retromer complex, causes late-onset Parkinson disease. Am. J. Hum. Genet. 89, 168–175 (2011).

13. Muhammad, A. et al. Retromer deficiency observed in Alzheimer’s disease causes hippocampal dysfunction, neurodegeneration, and Abeta accumulation. Proc. Natl. Acad. Sci. U. S. A. 105, 7327–32 (2008).

14. Wen, L. et al. VPS35 haploinsufficiency increases Alzheimer’s disease neuropathology. J. Cell Biol. 195, 765–779 (2011).

15. Vilariño-Güell, C. et al. VPS35 mutations in parkinson disease. Am. J. Hum. Genet. 89, 162–167 (2011).

16. Rovelet-Lecrux, A. et al. De novo deleterious genetic variations target a biological network centered on Aβ peptide in early-onset Alzheimer disease. Mol. Psychiatry 2015 209 20, 1046–1056 (2015).

17. Sargent, D. et al. Neuronal VPS35 deletion induces spinal cord motor neuron degeneration and early post-natal lethality. Brain Commun. 3, (2021).

18. Evans, A. J., Daly, J. L., Anuar, A. N. K., Simonetti, B. & Cullen, P. J. Acute inactivation of retromer and ESCPE-1 leads to time-resolved defects in endosomal cargo sorting. J. Cell Sci. 133, (2020).

19. Cataldo, A. M. et al. Endocytic pathway abnormalities precede amyloid β deposition in sporadic alzheimer’s disease and down syndrome: Differential effects of APOE genotype and presenilin mutations. Am. J. Pathol. 157, 277–286 (2000).

20. Crews, L. et al. Selective Molecular Alterations in the Autophagy Pathway in Patients with Lewy Body Disease and in Models of α-Synucleinopathy. PLoS One 5, e9313 (2010).

21. Shahmoradian, S. H. et al. Lewy pathology in Parkinson’s disease consists of crowded organelles and lipid membranes. Nat. Neurosci. 2019 227 22, 1099–1109 (2019).

22. Abu-Remaileh, M. et al. Lysosomal metabolomics reveals V-ATPase- and mTOR-dependent regulation of amino acid efflux from lysosomes. Science (80-.). 358, 807–813 (2017).

23. Curnock, R., Calcagni, A., Ballabio, A. & Cullen, P. J. TFEB controls retromer expression in response to nutrient availability. J. Cell Biol. 218, 3954–3966 (2019).

24. Pu, J. et al. BORC, a Multisubunit Complex that Regulates Lysosome Positioning. Dev. Cell 33, 176–188 (2015).

25. Bar-Peled, L. & Sabatini, D. M. Regulation of mTORC1 by amino acids. Trends Cell Biol. 24, 400–406 (2014).

26. Cui, Y. et al. Retromer has a selective function in cargo sorting via endosome transport carriers. J. Cell Biol. 218, 615– 631 (2019).

27. Gegg, M. E. & Schapira, A. H. V. The role of glucocerebrosidase in Parkinson disease pathogenesis. FEBS J. 285, 3591–3603 (2018).

28. Gieselmann, V. Lysosomal storage diseases. Biochim. Biophys. Acta - Mol. Basis Dis. 1270, 103–136 (1995).

29. Zerial, M. & McBride, H. Rab proteins as membrane organizers. Nat. Rev. Mol. Cell Biol. 2, 107–117 (2001).

30. Egami, Y. & Araki, N. Rab20 regulates phagosome maturation in RAW264 macrophages during Fc gamma receptor-mediated phagocytosis. PLoS One 7, (2012).

31. Jean, S., Cox, S., Nassari, S. & Kiger, A. A. Starvation-induced MTMR13 and RAB21 activity regulates VAMP8 to promote autophagosome–lysosome fusion. EMBO Rep. 16, 297 (2015).

32. Kasmapour, B., Gronow, A., Bleck, C. K. E., Hong, W. & Gutierrez, M. G. Size-dependent mechanism of cargo sorting during lysosome-phagosome fusion is controlled by Rab34. Proc. Natl. Acad. Sci. U. S. A. 109, 20485–20490 (2012).

33. Wilson, G. R. et al. Mutations in RAB39B Cause X-Linked Intellectual Disability and Early-Onset Parkinson Disease with α-Synuclein Pathology. Am. J. Hum. Genet. 95, 729–735 (2014).

34. Gonçalves, S. A. et al. shRNA-Based Screen Identifies Endocytic Recycling Pathway Components That Act as Genetic Modifiers of Alpha-Synuclein Aggregation, Secretion and Toxicity. PLOS Genet. 12, e1005995 (2016).

35. Hashimoto, Y., Shirane, M. & Nakayama, K. I. TMEM55B contributes to lysosomal homeostasis and amino acid– induced mTORC1 activation. Genes to Cells 23, 418–434 (2018).

36. Willett, R. et al. TFEB regulates lysosomal positioning by modulating TMEM55B expression and JIP4 recruitment to lysosomes. Nat. Commun. 2017 81 8, 1–17 (2017).

37. Dai, L. et al. Cholesterol Metabolism in Neurodegenerative Diseases: Molecular Mechanisms and Therapeutic Targets. Mol. Neurobiol. 2021 585 58, 2183–2201 (2021).

38. Graves, A. R., Curran, P. K., Smith, C. L. & Mindell, J. A. The Cl-/H+ antiporter ClC-7 is the primary chloride permeation pathway in lysosomes. Nat. 2008 4537196 453, 788–792 (2008).

39. Hu, M. et al. Parkinson’s disease-risk protein TMEM175 is a proton-activated proton channel in lysosomes. Cell 185, 2292–2308.e20 (2022).

40. O’Brien, R. J. & Wong, P. C. Amyloid precursor protein processing and Alzheimer’s disease. Annu. Rev. Neurosci. 34, 185–204 (2011).

41. Qureshi, Y. H. et al. The neuronal retromer can regulate both neuronal and microglial phenotypes of Alzheimer’s disease. Cell Rep. 38, 110262 (2022).

42. Nixon, R. A. Amyloid precursor protein and endosomal–lysosomal dysfunction in Alzheimer’s disease: inseparable partners in a multifactorial disease. FASEB J. 31, 2729 (2017).

43. Steinberg, F. et al. A global analysis of SNX27-retromer assembly and cargo specificity reveals a function in glucose and metal ion transport. Nat. Cell Biol. 15, 461–71 (2013).

44. Li, B. et al. The retromer complex safeguards against neural progenitor-derived tumorigenesis by regulating notch receptor trafficking. Elife 7, (2018).

45. Curnock, R. & Cullen, P. J. Mammalian copper homeostasis requires retromer-dependent recycling of the high-affinity copper transporter 1. J. Cell Sci. 133, (2020).

46. Kvainickas, A. et al. Retromer- and WASH-dependent sorting of nutrient transporters requires a multivalent interaction network with ANKRD50. J. Cell Sci. 130, 382–395 (2017).

47. Nakamura, F., Kalb, R. G. & Strittmatter, S. M. Molecular Basis of Semaphorin-Mediated Axon Guidance. J Neurobiol 44, 219–229 (2000).

48. Giniger, E. Notch signaling and neural connectivity. Curr. Opin. Genet. Dev. 22, 339 (2012).

49. Ivakine, E. A. et al. Neto2 is a KCC2 interacting protein required for neuronal Cl-regulation in hippocampal neurons. Proc. Natl. Acad. Sci. U. S. A. 110, 3561–3566 (2013).

50. Vernon, C. G. & Swanson, G. T. Neto2 Assembles with Kainate Receptors in DRG Neurons during Development and Modulates Neurite Outgrowth in Adult Sensory Neurons. J. Neurosci. 37, 3352 (2017).

51. Mahadevan, V. et al. Neto2-null mice have impaired GABAergic inhibition and are susceptible to seizures. Front. Cell. Neurosci. 9, 368 (2015).

52. del Puerto, A. et al. Kidins220 deficiency causes ventriculomegaly via SNX27-retromer-dependent AQP4 degradation. Mol. Psychiatry 2021 2611 26, 6411–6426 (2021).

53. Sebastián-Serrano, Á. et al. Differential regulation of Kidins220 isoforms in Huntington’s disease. Brain Pathol. 30, 120–136 (2020).

54. Li, Q. & Südhof, T. C. Cleavage of Amyloid-β Precursor Protein and Amyloid-β Precursor-like Protein by BACE 1. J. Biol. Chem. 279, 10542–10550 (2004).

55. Fotinopoulou, A. et al. BRI2 Interacts with Amyloid Precursor Protein (APP) and Regulates Amyloid β (Aβ) Production. J. Biol. Chem. 280, 30768–30772 (2005).

56. Matsuda, S. et al. The Familial Dementia BRI2 Gene Binds the Alzheimer Gene Amyloid-β Precursor Protein and Inhibits Amyloid-β Production. J. Biol. Chem. 280, 28912–28916 (2005).

57. Medina, D. L. et al. Transcriptional activation of lysosomal exocytosis promotes cellular clearance. Dev. Cell 21, 421– 30 (2011).

58. Alvarez-Erviti, L. et al. Lysosomal dysfunction increases exosome-mediated alpha-synuclein release and transmission. Neurobiol. Dis. 42, 360–367 (2011).

59. Miranda, A. M. et al. Neuronal lysosomal dysfunction releases exosomes harboring APP C-terminal fragments and unique lipid signatures. Nat. Commun. 9, 1–16 (2018).

60. Ballabio, A. & Bonifacino, J. S. Lysosomes as dynamic regulators of cell and organismal homeostasis. Nat. Rev. Mol. Cell Biol. 21, 101–118 (2020).

61. Follett, J. et al. The Vps35 D620N Mutation Linked to Parkinson’s Disease Disrupts the Cargo Sorting Function of Retromer. Traffic 15, 230–244 (2014).

62. Bugarcic, A. et al. Vps26A and Vps26B Subunits Define Distinct Retromer Complexes. Traffic 12, 1759–1773 (2011).

63. Rojas, R. et al. Regulation of retromer recruitment to endosomes by sequential action of Rab5 and Rab7. J. Cell Biol. 183, 513–26 (2008).

64. Hata, S. et al. Alcadein cleavages by amyloid β-precursor protein (APP) α- and γ-secretases generate small peptides, p3-Alcs, indicating Alzheimer disease-related γ-secretase dysfunction. J. Biol. Chem. 284, 36024–36033 (2009).

65. Walsh, R. B. et al. Opposing functions for retromer and Rab11 in extracellular vesicle traffic at presynaptic terminals. J. Cell Biol. 220, (2021).

66. Sardiello, M. et al. A gene network regulating lysosomal biogenesis and function. Science 325, 473–7 (2009).

67. Settembre, C. et al. A lysosome-to-nucleus signalling mechanism senses and regulates the lysosome via mTOR and TFEB. EMBO J. 31, 1095–108 (2012).

68. Palmieri, M. et al. Characterization of the CLEAR network reveals an integrated control of cellular clearance pathways. Hum. Mol. Genet. 20, 3852–66 (2011).

69. Peña-Llopis, S. et al. Regulation of TFEB and V-ATPases by mTORC1. EMBO J. 30, 3242–3258 (2011).

70. Mansueto, G. et al. Transcription Factor EB Controls Metabolic Flexibility during Exercise. Cell Metab. 25, 182–196 (2017).

71. Seaman, M. N. J. Identification of a novel conserved sorting motif required for retromer-mediated endosome-to-TGN retrieval. J. Cell Sci. 120, 2378–2389 (2007).

72. Lucas, M. et al. Structural Mechanism for Cargo Recognition by the Retromer Complex. Cell 167, 1623–1635.e14 (2016).

73. Ostrowski, M. et al. Rab27a and Rab27b control different steps of the exosome secretion pathway. Nat. Cell Biol. 12, 19–30 (2010).

74. Joshi, C. S., Mora, A., Felder, P. A. & Mysorekar, I. U. NRF2 promotes urothelial cell response to bacterial infection by regulating reactive oxygen species and RAB27B expression. Cell Rep. 37, (2021).

75. Underwood, R. et al. The GTPase Rab27b regulates the release, autophagic clearance, and toxicity of alpha-synuclein. J. Biol. Chem. 295, 8005–8016 (2020).

76. Underwood, R., Wang, B., Pathak, A., Volpicelli-Daley, L. & Yacoubian, T. A. Rab27 GTPases regulate alpha-synuclein uptake, cell-to-cell transmission, and toxicity. bioRxiv 2020.11.17.387449 (2020) doi:10.1101/2020.11.17.387449.

77. Jia, D. et al. Structural and mechanistic insights into regulation of the retromer coat by TBC1d5. Nat. Commun. 7, 1– 11 (2016).

78. Hesketh, G. et al. VARP is recruited on to endosomes by direct interaction with retromer, where together they function in export to the cell surface. Dev. Cell 29, 591–606 (2014).

79. Yu, L. et al. Termination of autophagy and reformation of lysosomes regulated by mTOR. Nature 465, 942–946 (2010).

80. Wu, K. et al. BLOC1S1/GCN5L1/BORCS1 is a critical mediator for the initiation of autolysosomal tubulation. Autophagy 17, 3707–3724 (2021).

81. McGrath, M. J. et al. Defective lysosome reformation during autophagy causes skeletal muscle disease. J. Clin. Invest. 131, (2021).

82. Mecozzi, V. J. et al. Pharmacological chaperones stabilize retromer to limit APP processing. Nat. Chem. Biol. 10, 443– 9 (2014).

83. Muzio, L. et al. Retromer stabilization results in neuroprotection in a model of Amyotrophic Lateral Sclerosis. Nat. Commun. 11, 1–17 (2020).

84. Luzio, J. P., Pryor, P. R. & Bright, N. A. Lysosomes: Fusion and function. Nat. Rev. Mol. Cell Biol. 8, 622–632 (2007).

85. Chen, Y. & Yu, L. Recent progress in autophagic lysosome reformation. Traffic 18, 358–361 (2017).

86. Zavodszky, E. et al. Mutation in VPS35 associated with Parkinson’s disease impairs WASH complex association and inhibits autophagy. Nat. Commun. 5, 3828 (2014).

87. Pu, J., Keren-Kaplan, T. & Bonifacino, J. S. A Ragulator-BORC interaction controls lysosome positioning in response to amino acid availability. J. Cell Biol. 216, 4183–4197 (2017).

88. Yordanov, T. E. et al. Biogenesis of lysosome-related organelles complex-1 (BORC) regulates late endosomal/lysosomal size through PIKfyve-dependent phosphatidylinositol-3,5-bisphosphate. Traffic 20, 674–696 (2019).

89. Rong, Y. et al. Clathrin and phosphatidylinositol-4,5-bisphosphate regulate autophagic lysosome reformation. Nat. Cell Biol. 14, 924–934 (2012).

90. Du, W. et al. Kinesin 1 Drives Autolysosome Tubulation. Dev. Cell 37, 326–336 (2016).

91. Dai, A., Yu, L. & Wang, H. W. WHAMM initiates autolysosome tubulation by promoting actin polymerization on autolysosomes. Nat. Commun. 10, (2019).

92. Bécot, A., Volgers, C. & van Niel, G. Transmissible endosomal intoxication: A balance between exosomes and lysosomes at the basis of intercellular amyloid propagation. Biomedicines 8, (2020).

93. Simonetti, B., Danson, C. M., Heesom, K. J. & Cullen, P. J. Sequence-dependent cargo recognition by SNX-BARs mediates retromer-independent transport of CI-MPR. J. Cell Biol. 216, 3695–3712 (2017).

94. Costes, S. V. et al. Automatic and quantitative measurement of protein-protein colocalization in live cells. Biophys. J. 86, 3993–4003 (2004).

95. Starling, G. P. et al. Folliculin directs the formation of a Rab34– RILP complex to control the nutrient-dependent dynamic distribution of lysosomes. EMBO Rep. 17, 823–841 (2016).

96. Xiong, Y. et al. A Comparison of mRNA Sequencing with Random Primed and 3′-Directed Libraries. Sci. Reports 2017 71 7, 1–12 (2017).

97. Dobin, A. et al. STAR: ultrafast universal RNA-seq aligner. Bioinformatics 29, 15–21 (2013).

98. Anders, S., Pyl, P. T. & Huber, W. HTSeq—a Python framework to work with high-throughput sequencing data. Bioinformatics 31, 166–169 (2015).

99. Robinson, M. D., McCarthy, D. J. & Smyth, G. K. edgeR: a Bioconductor package for differential expression analysis of digital gene expression data. Bioinformatics 26, 139–140 (2010).

100. Mootha, V. K. et al. PGC-1α-responsive genes involved in oxidative phosphorylation are coordinately downregulated in human diabetes. Nat. Genet. 34, 267–273 (2003).

101. Subramanian, A. et al. Gene set enrichment analysis: A knowledge-based approach for interpreting genome-wide expression profiles. Proc. Natl. Acad. Sci. U. S. A. 102, 15545–15550 (2005).

102. Merico, D., Isserlin, R., Stueker, O., Emili, A. & Bader, G. D. Enrichment map: A network-based method for gene-set enrichment visualization and interpretation. PLoS One 5, (2010).

103. Durinck, S., Spellman, P. T., Birney, E. & Huber, W. Mapping identifiers for the integration of genomic datasets with the R/Bioconductor package biomaRt. Nat. Protoc. 2009 48 4, 1184–1191 (2009).

104. Goedhart, J. & Luijsterburg, M. S. VolcaNoseR is a web app for creating, exploring, labeling and sharing volcano plots. Sci. Rep. 10, 1–5 (2020).

105. Zhou, Y. et al. Metascape provides a biologist-oriented resource for the analysis of systems-level datasets. Nat. Commun. 2019 101 10, 1–10 (2019).

106. Mi, H. et al. Protocol Update for large-scale genome and gene function analysis with the PANTHER classification system (v.14.0). Nat. Protoc. 14, 703–721 (2019).

